# A Cell Atlas of Microbe-Responsive Processes in the Zebrafish Intestine

**DOI:** 10.1101/2020.11.06.371609

**Authors:** Reegan J. Willms, Lena Ocampo Jones, Jennifer C. Hocking, Edan Foley

**Affiliations:** Department of Medical Microbiology and Immunology, Faculty of Medicine and Dentistry, University of Alberta, Edmonton, AB, Canada; Division of Anatomy, Department of Surgery, Faculty of Medicine and Dentistry, University of Alberta, Edmonton, AB, Canada

## Abstract

Gut microbial products direct growth, differentiation, and development in the animal host. Disruptions to host-microbe interactions have profound health consequences, that include onset of chronic inflammatory illnesses. However, we lack system-wide understanding of cell-specific responses to the microbiome. We profiled transcriptional activity in individual cells from the intestine, and associated tissue, of zebrafish larvae that we raised in the presence or absence of a microbiome. We uncovered extensive cellular heterogeneity in the conventional zebrafish intestinal epithelium, including previously undescribed cell types with known mammalian homologs. By comparing conventional to germ-free profiles, we mapped microbial impacts on transcriptional activity in each cell population. We revealed intricate degrees of cellular specificity in host responses to the microbiome that included regulatory effects on patterning, metabolic and immune activity. For example, we showed that removal of microbes hindered pro-angiogenic signals in the developing vasculature, resulting in impaired intestinal vascularization. Our work provides a high-resolution atlas of intestinal cellular composition in the developing fish gut and details the effects of the microbiome on each cell type. Furthermore, we provide a web-based resource for single-cell gene expression visualization under conventional and germ-free conditions to facilitate exploration of this dataset.

## INTRODUCTION

Research conducted with a variety of model organisms has revealed much about the importance of gut microbes for host health. Animals raised in sterile, germ-free environments frequently exhibit defects in growth, immunity and metabolism (Bates et al., 2006; Hooper et al., 2001; Rawls et al., 2004; Reikvam et al., 2011). Of equal importance, changes in composition or distribution of gut microbial communities are associated with severe and sometimes deadly illnesses, including gastrointestinal cancers and inflammatory bowel diseases (Belkaid and Hand, 2014; Zitvogel et al., 2015). Thus, it is critical to fully understand how the microbiota impacts development, growth, and cellular function of host organisms.

Zebrafish larvae have emerged as a valuable tool to identify key regulators of host-microbe interactions (Brugman, 2016; Flores et al., 2020; López Nadal et al., 2020). Zebrafish embryos develop within a protective chorion that shields them from environmental microbes up to forty-eight hours post fertilization (hpf). Once larvae exit the chorion, water-borne microbes colonize the gut lumen (Bates et al., 2006; Stephens et al., 2016; Wallace et al., 2005), where they influence host development (Bates et al., 2006; Cheesman et al., 2011; Kanther et al., 2011; Koch et al., 2018). From a technical perspective, zebrafish offer several advantages to pinpoint developmental responses to the microbiota. Larvae are amenable to sophisticated manipulations, including genetic modifications from the single cell stage (Grunwald and Eisen, 2002). Additionally, researchers have simple protocols to generate large numbers of germ-free larvae, or larvae associated with defined microbial communities (Melancon et al., 2017; Pham et al., 2008), and the translucent epidermis is ideal for visualization of internal structures in fixed or live samples. Thus, zebrafish provide a convenient window to visualize microbial controls of vertebrate physiology.

Importantly, genetic regulation of intestinal function is highly similar between zebrafish and mammals. In both systems, orthologous signals, including those driven by Notch, Bone Morphogenetic Protein (BMP) and Wnt pathways, direct development of absorptive and secretory cell lineages from cycling progenitors in the intestinal epithelium (Cheesman et al., 2011; Crosnier et al., 2005; Davison et al., 2017; Flasse et al., 2013; Haramis et al., 2006; Muncan et al., 2007; Roach et al., 2013; Yang et al., 2009). While the zebrafish intestine possesses mucin-producing goblet cells and regulatory enteroendocrine cells (Crosnier et al., 2005; Ng et al., 2005; Wallace et al., 2005), there are no reports of immune-modulatory Paneth or tuft cells. Our understanding of absorptive lineages is less complete. Like mammals, the fish intestinal epithelium includes regionally specialized enterocytes that harvest nutrients from the lumen (Lickwar et al., 2017; Ng et al., 2005; Park et al., 2019; Wallace et al., 2005; Wang et al., 2010b). However, the extent of functional heterogeneity within enterocyte populations is unclear, and we do not know if the intestinal epithelium houses specialized absorptive cells such as antigen-capturing M cells, or recently described Best4/Otop2 cells (Parikh et al., 2019; Smillie et al., 2019). Likewise, despite experimental evidence for the existence of cycling progenitors (Crosnier et al., 2005; Li et al., 2020; Peron et al., 2020; Rawls et al., 2004; Wallace et al., 2005), we lack expression markers that permit identification and manipulation of this essential cell type. Combined, these deficits have hampered our ability to harness the full potential of the zebrafish as a model of intestinal biology and host-microbe interactions.

From a microbial perspective, similarities between fish and mammals are also evident. Like mammals, fish rely on a complex network of germline-encoded innate defenses, and lymphocyte-based adaptive defenses to prevent invasion of interstitial tissues by gut-resident microbes (Flores et al., 2020; Hernández et al., 2018). Transcriptional studies showed that orthologous genes mediate microbe-dependent control of epithelial proliferation, nutrient metabolism, xenobiotic metabolism, and innate immunity (Davison et al., 2017; Heppert et al., 2021; Hooper et al., 2001; Koch et al., 2018; Rawls et al., 2004; Reikvam et al., 2011). *In vivo* studies support a shared role for the microbiota in developmental processes including epithelial renewal, secretory cell differentiation, and gut motility (Cheesman et al., 2011; Troll et al., 2018; Wiles et al., 2016). Microbes also educate immune systems in fish and mammals, inducing mucosal inflammation and myeloid cell recruitment through Myd88-dependent TLR signals (Galindo-Villegas et al., 2012; Koch et al., 2018; Takeda and Akira, 2005). Like mammals, fish neutralize pathogenic bacteria via epithelial production of reactive oxygen species and antimicrobial peptides (Flores et al., 2010; Katzenback, 2015). Additionally, experimental evidence in fish revealed that epithelial alkaline phosphatase detoxifies LPS, a finding later corroborated in mice (Bates et al., 2007; Goldberg et al., 2008). These findings demonstrate that zebrafish, alongside other models, can inform our understanding of microbial impacts on host development and disease.

While recent studies provide significant insights into microbe-dependent host processes, much of this work focused on how microbes impact whole organisms, the entire intestine, or intestinal regions. Thus, a knowledge gap exists in our understanding of cell type-specific processes reliant on microbial signals. A few studies addressed this disparity via fluorescent activated cell sorting of intestinal epithelial sub-populations (Arora et al., 2018), however this method requires foreknowledge of, and access to, cell type-specific antibodies or reporters, and assays are limited to one cell type per experiment.

To achieve unbiased cell type-specific analysis of microbe-dependent processes, and to advance cellular characterization of the fish intestine, we prepared single cell transcriptional atlases of intestines from 6 days post fertilization (dpf) zebrafish larvae raised in a conventional environment, or in the absence of a microbiome. We identified thirty-five distinct transcriptional states in the intestine, several of which were previously undescribed, and completed a high-resolution map of cellular responses to the microbiota that showed cell-specific microbial effects on growth, patterning, and immunity in the host. To facilitate community-wide mining of our results, we have made both sets publicly accessible for user-friendly visualization on the Broad Institute Single Cell Portal

## RESULTS

### A single cell atlas of the zebrafish larval intestine

To trace effects of commensal microbes on intestinal physiology, we prepared single-cell transcriptional profiles of digestive tracts from 6 dpf zebrafish larvae raised under conventional (CV) or germ-free (GF) conditions (Fig. 1A and Supplementary Fig. 1). Our results included tissues that exist in close association with the gut, such as the pancreas and liver. After filtering for dead cells and doublets, we determined gene expression profiles for 18,345 individual cells (8,036 CV; 10,309 GF; Fig. 1B). To advance our understanding of cellular heterogeneity within the intestine, we used graph-based clustering to identify cell types within our integrated data (Fig. 1B). We identified 35 distinct clusters (Supplementary Table 1), which we grouped into 18 cell types based on expression of known markers (Fig. 1B).

**Figure 1.**
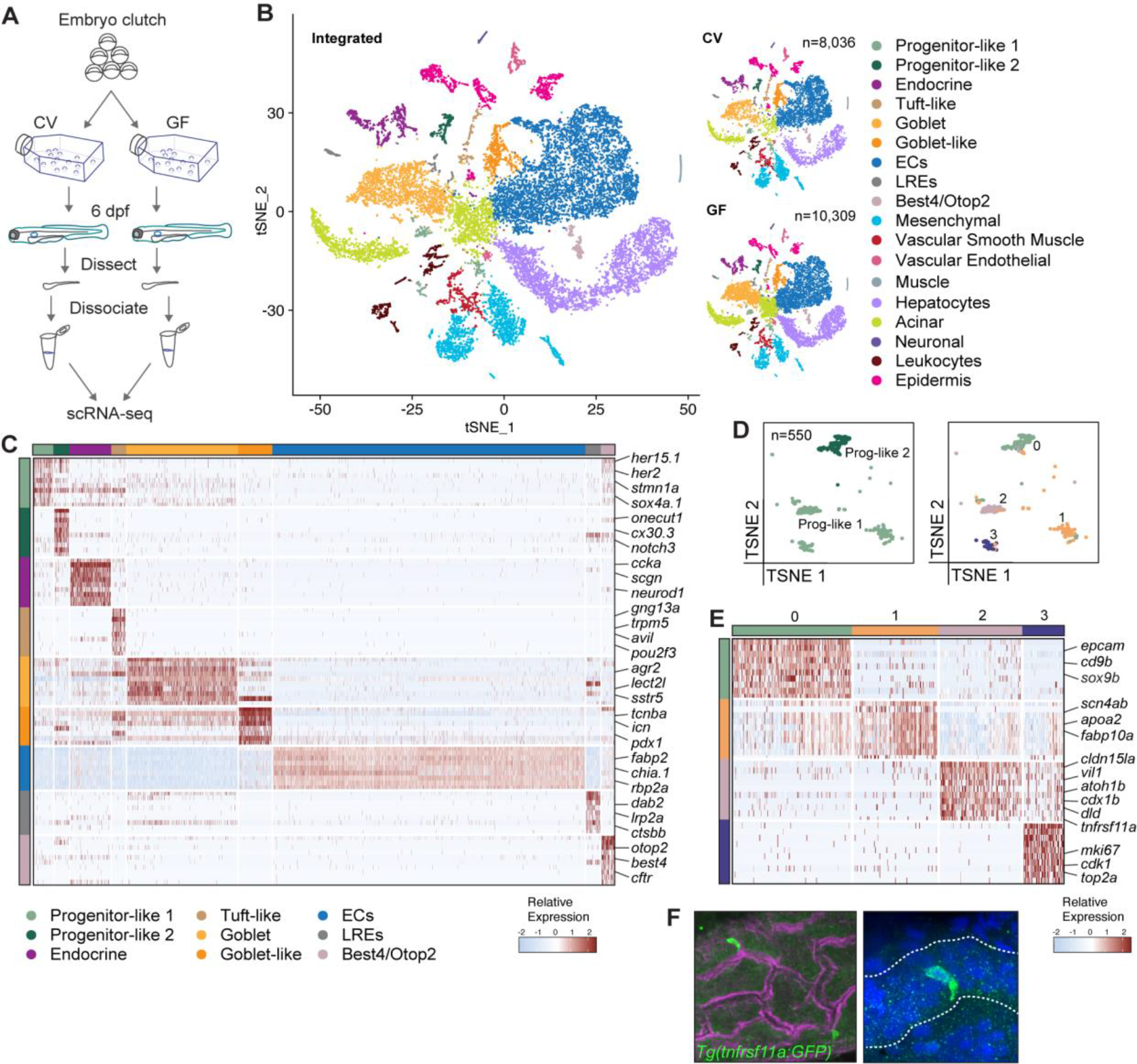
Transcriptionally distinct cell populations in the zebrafish intestine. (*A*) Experimental design for transcriptional profiling of single cells in the zebrafish intestine. (*B*) 2D t-SNE projections of profiled cells color coded by cell type. Left panel shows t-SNE of integrated CV and GF datasets, with CV and GF conditions shown independently on the top and bottom right respectively. (*C*) Heatmap of IEC cluster markers colored by relative gene expression. Cell types are indicated by colored bars on the left and top. Several top markers for each cluster are shown on the right axis of the heatmap. (*D*) t-SNE plots of progenitor-like clusters 1 and 2 from original graph-based analysis (left) and further re-clustering (right), color coded by cell type. (*E*) Heatmap of cell markers for putative progenitor-like clusters, colored by relative gene expression. Cell types are indicated by colored bars on the left and top. Several top markers for each cluster are shown on the right axis of the heatmap. (*F*) Optical section of a whole gut from 6 dpf *Tg(tnfrsf11a:GFP)* zebrafish, stained with phalloidin to visualize filamentous actin (magenta) and Hoechst to visualize nuclei (blue). Right panel is a magnified image of a GFP positive cell from the left panel.

Our datasets were dominated by expression profiles for intestinal epithelial cells (IECs). For example, we identified secretory peptide hormone-producing enteroendocrine cells, as well as goblet cells marked by expression of the goblet cell differentiation factor *anterior gradient 2* (*agr2*), and *sstr5* (Fig. 1C), a gene product that stimulates Mucin 2 production in the mouse colon (Song et al., 2020). We also uncovered a goblet-like cluster that upregulated *pdx1* (Fig. 1C), enriched in secretory cells of the foregut and pancreas (Lavergne et al., 2020). Besides endocrine and goblet cell lineages, we identified an unexpected cluster with pronounced transcriptional similarity to mammalian intestinal tuft cells, including expression of tuft cell marker genes *Gng13, Trpm5, Avil*, and the tuft cell specification master regulator *Pou2f3* (Haber et al., 2017) (Fig. 1C and Supplementary Fig. 2A-C). Transmission electron microscopy of adult zebrafish intestinal epithelia uncovered a rare, rotund cell type with classical morphological features of intestinal tuft cells (Hoover et al., 2017), namely a tubular cytoskeletal network below an apical tuft of microvilli (Supplementary Fig. 2D). Thus, our data suggest that, like mammals, zebrafish may contain a rare population of sensory intestinal tuft cells.

The majority of IECs were absorptive cells and included canonical enterocyte (EC) lineages that expressed genes required for nutrient acquisition and metabolism, as well as recently described lysosome-rich enterocytes (LREs) (Fig. 1C and Supplementary Table 1), thought to mediate protein degradation (Park et al., 2019). Separately, we discovered a population of Best4/Otop2 cells (Fig. 1B-C and Supplementary Table 1), an absorptive lineage recently described in humans (Parikh et al., 2019; Smillie et al., 2019), and uncharacterized in zebrafish. Like human Best4/Otop2 cells, the fish counterparts were marked by enhanced expression of *notch2* and Notch-responsive *hes-related* family members (Fig. 1C, Supplementary Table 1 and Supplementary Fig. 3). Additionally, zebrafish Best4/Otop2 cells expressed the chloride/bicarbonate transporter *cftr* (Fig. 1C and Supplementary Table 1), suggesting possible functional similarities with human duodenal BCHE cells (Busslinger et al., 2021).

Apart from absorptive and secretory lineages, our initial clustering uncovered two populations that displayed features associated with intestinal progenitor cells, including expression of Notch pathway components *dld*, *dla,* and *HES5* othologues *her15* and *her2* (progenitor-like 1), as well as *notch3* (progenitor-like 2) (Fig 1C and Supplementary Table 1). A more detailed analysis resolved the putative progenitor pool into four sub-clusters with distinct transcriptional hallmarks (Fig. 1D-E). Of these four, we believe cluster one is hepatic in origin, as it is marked by expression of liver-associated genes *apoa2* and *fabp10a* (Fig. 1E). In contrast, cells from clusters zero, two, and three had features frequently associated with intestinal progenitors. Cluster zero was marked by expression of the gut-associated genes *onecut1* (Matthews et al., 2004) and *notch3* (Crosnier et al., 2005) in addition to *sox9b* (Fig. 1E), an intestinal stem cell marker in medaka fish (Aghaallaei et al., 2016), and a marker of basal columnar IECs in adult zebrafish (Peron et al., 2020). Additionally, cluster zero cells expressed elevated amounts of *epcam* (Fig. 1E), a gene linked with intestinal epithelial proliferation in vertebrates (Ouchi et al., 2021). Cluster two cells expressed intestinal epithelial cell markers *cldn15la* (Alvers et al., 2014) and *vil1* (Abrams et al., 2012; Thakur et al., 2014), as well as regulators of intestinal progenitor cell division and differentiation, such as *cdx1b* (Flores et al., 2008) and *atoh1b* (Fig. 1E). Furthermore, fluorescence imaging of intestines from *Tg(tnfrsf11a:GFP)* fish that expressed GFP under control of the promoter for cluster two marker *tnfrsf11a* showed that, like intestinal progenitors (Li et al., 2020; Ng et al., 2005), cluster two cells reside at the base of intestinal folds (Fig. 1F). Finally, we identified cluster three as a cycling population that actively expressed proliferation markers of mammalian transit amplifying cells (Haber et al., 2017), such as *mki67*, *cdk1*, and *top2a* (Fig. 1E). Thus, our transcriptional and *in vivo* data identified a previously undescribed pool of IECs with hallmarks of intestinal progenitors, although lineage tracing studies are required for confirmation. In sum, we have identified a panel of expression markers that distinguish major lineages of the zebrafish digestive tract, including previously undescribed tuft-like cells, Best4/Otop2 cells, and possible markers of intestinal progenitors.

### Cell type-specific effects of gut microbes on host gene expression

Despite critical roles for microbial factors in regulation of host physiology, we have made sporadic progress charting cell type-specific responses to the microbiome. Like other facilities (Roeselers et al., 2011), our fish primarily host γ- and α-Proteobacteria (Supplementary Tables 2 and 3). Therefore, we believe that comparisons between our GF and CV data may uncover relevant cell-specific responses to the microbiome.

We first confirmed that our data reproduce known effects of GF growth on host gene expression. Among the top globally differentially expressed genes in GF fish relative to CV controls, we identified known microbe-dependent effects on expression of a range of host genes (Rawls et al., 2004), including *gpx1b*, *socs3a*, and *tyrosine aminotransferase* (*tat*) (Fig. 2A). Additionally, we used Nanostring quantitative analysis to independently validate effects of the microbiome on expression of host genes observed in our single cell data (Fig. 2B). Finally, of 175 microbe-responsive genes identified by Rawls and co-authors (2004), we observed 125 (71%) with significant microbe-dependent expression changes in at least one cell subset (Supplementary Table 4). Collectively, these observations argue that our gene expression data accurately report effects of the microbiome on gut function.

**Figure 2.**
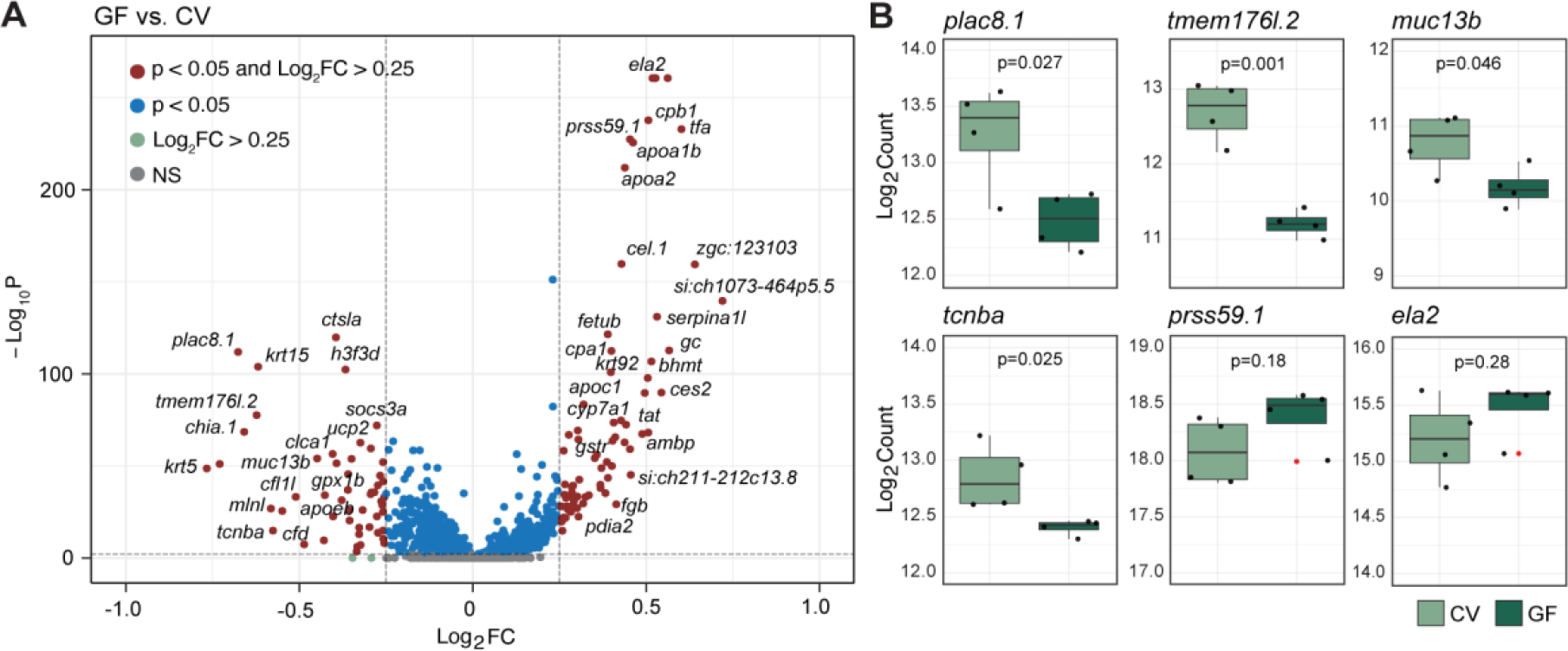
Microbial control of host gene expression. (*A*) Volcano plot of differentially expressed genes in GF relative to CV cells, treated in aggregate. Significance was determined in Seurat using the non- parametric Wilcoxon rank sum test. (*B*) Boxplots of Nanostring gene expression analysis from dissected whole guts. Four replicates (n=15 guts per replicate) were analyzed per condition. Outliers are indicated with red dots. Significance was determined using a Student’s *t*-test.

In some cases, such as *tat*, elimination of the microbiome altered gene expression throughout the gut (Supplementary Table 4). However, we also observed instances where GF growth impacted gene expression in specific cell types. For example, removal of the microbiome attenuated *moesin a* (*msna*) expression exclusively in vascular endothelial cells, vascular smooth muscle, progenitors, and leukocytes (Supplementary Table 4). To explore cell-type specific microbiome responses in greater detail, we characterized the transcriptional programs of progenitor-like cells raised under CV and GF conditions. We selected progenitors, as microbes are established modifiers of proliferation and differentiation, including Notch pathway components (Crosnier et al., 2005; Flasse et al., 2013; Roach et al., 2013; Yang et al., 2009). We observed remarkable cellular specificity in the responses of putative progenitor clusters to GF growth (Supplementary Fig. 4). For example, cells from progenitor-like cluster two downregulated Notch-responsive transcription factors *atoh1b* and *her15.1*, as well as the intestinal Notch ligand *delta D* (*dld*) when grown in the absence of a microbiome (Supplementary Fig. 4B-D). Intriguingly, we also observed decreased expression of *interferon-related developmental regulator 1* (*ifrd1*) in progenitor-like cluster two (Supplementary Fig. 4D), an immune response gene that regulates gut epithelial proliferation (Yu et al., 2010). Thus, our data indicate that a specific subset of candidate progenitors are particularly sensitive to the impacts of microbial factors on Notch activity.

To test the utility of our data for cell-specific mapping of signaling pathway activity in the presence or absence of microbes, we visualized relative expression of microbial sensors, NF-kB pathway components, cytokines and chemokines in CV and GF fish. In CV larvae, we detected cell-restricted expression of key immune sensors and effectors (Supplementary Fig. 5). For example, leukocytes expressed immune-regulatory cytokines such as *cxcl8a*, *il1b,* and *tnfa*, whereas the vasculature was characterized by enriched expression of microbial sensor *tlr4ba*, cytokine *tgfb1b* and the inflammation regulator *ahr2*. Like mammals, CV hepatocytes expressed the *hamp* antimicrobial peptide, while mesenchymal cells were characterized by enriched expression of *cxcl8b* isoforms, *cxcl12*, and the *tgfb1a*, *tgfb2* and *tgfb3* cytokines. Within the intestinal epithelium, most enterocyte subtypes produced *alpi.2*, a phosphatase required for detoxification of bacterial lipopolysaccharide (Bates et al., 2007), whereas enteroendocrine cells expressed *il22*, a cytokine that activates epithelial innate defenses (Dudakov et al., 2015). In agreement with previous reports (Kanther et al., 2011), *serum amyloid A* (*saa*) was expressed in mid-intestinal LREs and goblet cells. Intriguingly, we also saw enhanced *nod1, nod2* and *myd88* expression in CV tuft-like cells, consistent with proposed roles for Nod1 and Nod2 in type 2 immunity in tuft cells (Magalhaes et al., 2011). Removal of the microbiome significantly impacted organization of immune pathways in developing larvae. In particular, we noted greatly diminished expression of *il22* from endocrine cells and leukocytes; *nod1* from progenitors, mesenchyme, and vasculature; *ifit14* and *ifit15* (*IFIT1* orthologues) from enterocytes; *stat2* across IECs; and *tgf* isoforms from mesenchymal cells, vasculature, and leukocytes. Combined, our data uncover a sophisticated partitioning of immune gene expression patterns across CV cell types, many of which indicate shared microbe-response pathways in zebrafish and mammals. Importantly, we believe the utility of our gene expression data extends beyond identification of cell-specific immune activity. To facilitate community-wide mining of our results for all pathways or genes of interest, we have made both sets publicly available for user-friendly visualization on the Broad Institute Single Cell Portal.

### Microbes regulate larval secretory lineage functions

Zebrafish secretory lineages primarily consist of hormone-producing enteroendocrine cells and mucus-secreting goblet cells. To understand how secretory cells interact with a conventional microbiome, we generated a transcriptional atlas of secretory cells from CV and GF fish. Among the enteroendocrine population, we uncovered six distinct transcriptional cell states, each marked by a unique pattern of peptide hormone production (Fig. 3A-B) and distinct spatial distribution profiles (Supplementary Fig. 6A). For example, CV enteroendocrine cluster five cells expressed anterior intestinal markers, and were characterized by production of *ccka* and *cckb*, regulators of gut motility, satiety, and lipid and protein digestion (Le et al., 2019; Rehfeld, 2017). By contrast, enteroendocrine cluster three cells appeared to have a more diffuse rostro-caudal distribution and were the predominant source of the motility regulator *vipb*, and the multifunctional peptide *galn*. We observed modest effects of GF growth on expression of most peptide hormones, suggesting that enteroendocrine lineage specification is broadly insensitive to microbial exposure. However, we detected instances where microbial presence significantly affected hormone expression profiles of distinct enteroendocrine lineages. In particular, we observed significantly diminished expression of *gip* and *gcgb* within cluster four enteroendocrine cells, as well as enhanced expression of the appetite suppressant *pyyb* in cluster five cells. These data support roles for microbes in modifying levels of *gip* and *glucagon*, incretin hormones that regulate glucose metabolism and insulin secretion (Gribble and Reimann, 2016), and further implicate microbes in the control of *pyyb* production. Upon examination of goblet cells, we identified two clusters (Fig. 3C) defined by highly similar gene expression profiles (Supplementary Table 1), where cluster one was enriched for *agr2* (Supplementary Fig. 6B), and both clusters primarily expressed mid-intestinal markers (Supplementary Fig. 6A), consistent with goblet cell distribution in zebrafish guts (Ng et al., 2005; Wallace et al., 2005). Furthermore, we identified an *agr2*-enriched goblet-like cluster that expressed *mucin 5.3* (Supplementary Fig. 6C), known to be enriched in the esophagus (Jevtov et al., 2014), suggesting that this cluster may represent mucus-producing foregut cells.

**Figure 3.**
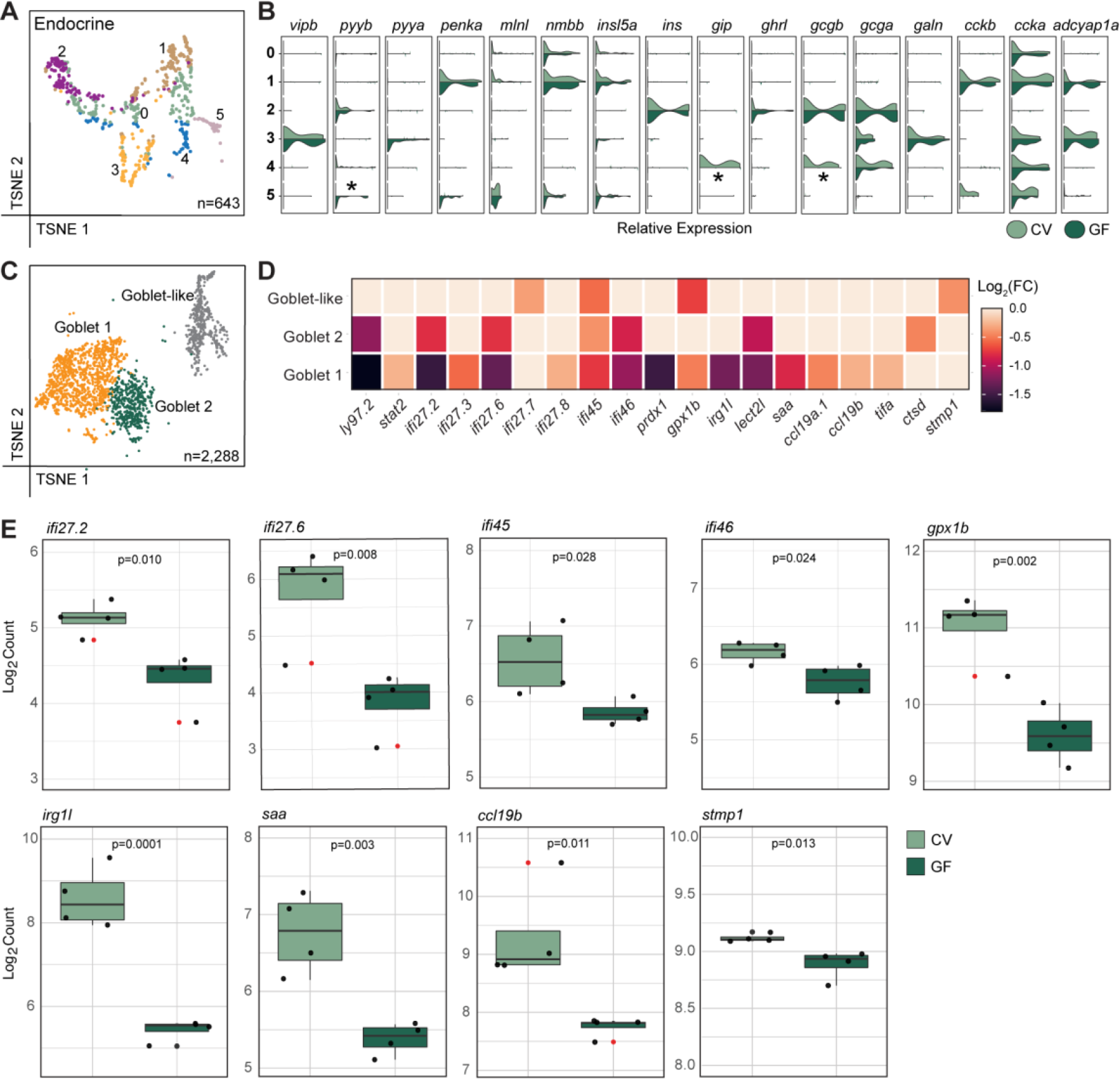
Germ-free growth alters peptide hormone expression in enteroendocrine cells and immune signaling in goblet cells. (*A*) t-SNE plot of enteroendocrine cells after re-clustering, color coded by cell type. (*B*) Violin plots for expression of zebrafish peptide hormones, as expressed in enteroendocrine clusters 0-5. Asterisks indicate significant gene expression differences (p<0.05) between GF and CV conditions, as determined with a non-parametric Wilcoxon rank sum test. (*C*) t-SNE plot of goblet and goblet-like cell clusters color coded by cell type. (*D*) Heatmap of differentially expressed immune related genes in GF relative to CV cell populations, color coded according to Log2(FC). All non-zero value expression changes are significant (p<0.05) as determined with a non-parametric Wilcoxon rank sum test. (*E*) Boxplots of Nanostring gene expression analysis from dissected whole guts. Four replicates (n=15 guts per replicate) were analyzed per condition. Outliers are indicated with red dots. Significance was determined using a Student’s *t*-test.

We were intrigued by apparent changes to immunity in GF goblet cells relative to CV counterparts (Supplementary Fig. 5), so we examined goblet cell immune gene expression in greater detail. Removal of the microbiome had cluster-specific impacts on several immune regulators. For example, microbiome elimination resulted in significantly diminished expression of the putative LPS-binding molecule and anti- microbial peptide *ly97.2* (Liu et al., 2017; Wang et al., 2016), as well as the inflammation mediators *irg1l* (Hall et al., 2014; van Soest et al., 2011) and *lect2l* (Gonçalves et al., 2012) in goblet cell clusters 1 and 2. In contrast, GF growth led to diminished expression of *interferon alpha inducible protein* 27 (*IFI27*) orthologues across all goblet cells, whereas absence of the microbiome exclusively attenuated expression of *CCL19* orthologues in cluster one cells (Fig. 3D). We validated microbiome-dependent expression changes to several genes including *IFI27* orthologues, *irg1l*, *ccl19b*, and IL-1β regulator *stmp1* by whole-tissue Nanostring gene expression analysis (Fig. 3E), further supporting a role for goblet cell-mediated IFN and inflammatory signaling in response to commensal microbes. In short, our data uncovered a complex arrangement of goblet and enteroendocrine IECs with non-overlapping rostro-caudal distribution and subset-specific responses to a conventional microbiome.

### The zebrafish intestinal epithelium houses functionally diverse absorptive lineages

Like most animals, the zebrafish intestinal epithelium primarily contains absorptive cells that acquire material from the gut lumen. In fish, metabolite acquisition relies on specialist enterocyte and protein- acquiring LRE lineages (Lickwar et al., 2017; Ng et al., 2005; Park et al., 2019; Wallace et al., 2005; Wang et al., 2010b). To characterize microbial responsiveness and functional specializations within each lineage, we analyzed gene expression in absorptive clusters from our integrated CV and GF datasets. We identified five enterocyte clusters (Fig. 4A), of which clusters one to four were enriched in the anterior intestine (Fig. 4D) and marked by expression of genes required for lipid, carbohydrate, chitin, and small molecule metabolism (Fig. 4B-C and Supplementary Table 1). Cluster five cells were a distinct subset, specialized in the metabolism of xenobiotic compounds (Fig. 4C), and transport of vitamin B12 by *transcobalamin beta a* (*tcnba*) (Fig. 4B and Supplementary Table 1). Alongside enterocytes, we captured expression profiles for three separate absorptive lineages, two of which had expression profiles consistent with LREs (Park et al., 2019). LRE1 cells were relatively rare and expressed pronephros markers such as *lrp2a*, *zgc:64022* and *tspan35* (Fig. 4B and Supplementary Table 1), suggesting that LRE1 cells are renal. In contrast, LRE2 cells appear mid-intestinal (Fig. 4D), enriched for expression of genes required for peptide catabolism, in agreement with a role for mid-intestinal LREs in protein digestion (Park et al., 2019). Lastly, we identified a previously unknown absorptive lineage analogous to recently characterized human colonic BEST4/OTOP2 cells (Parikh et al., 2019; Smillie et al., 2019). Like the human equivalent, zebrafish Best4/Otop2 cells were a posterior cell type (Fig. 4D), that expressed genes required for ion transport (*cftr*, *ca2*, *best4*).

**Figure 4.**
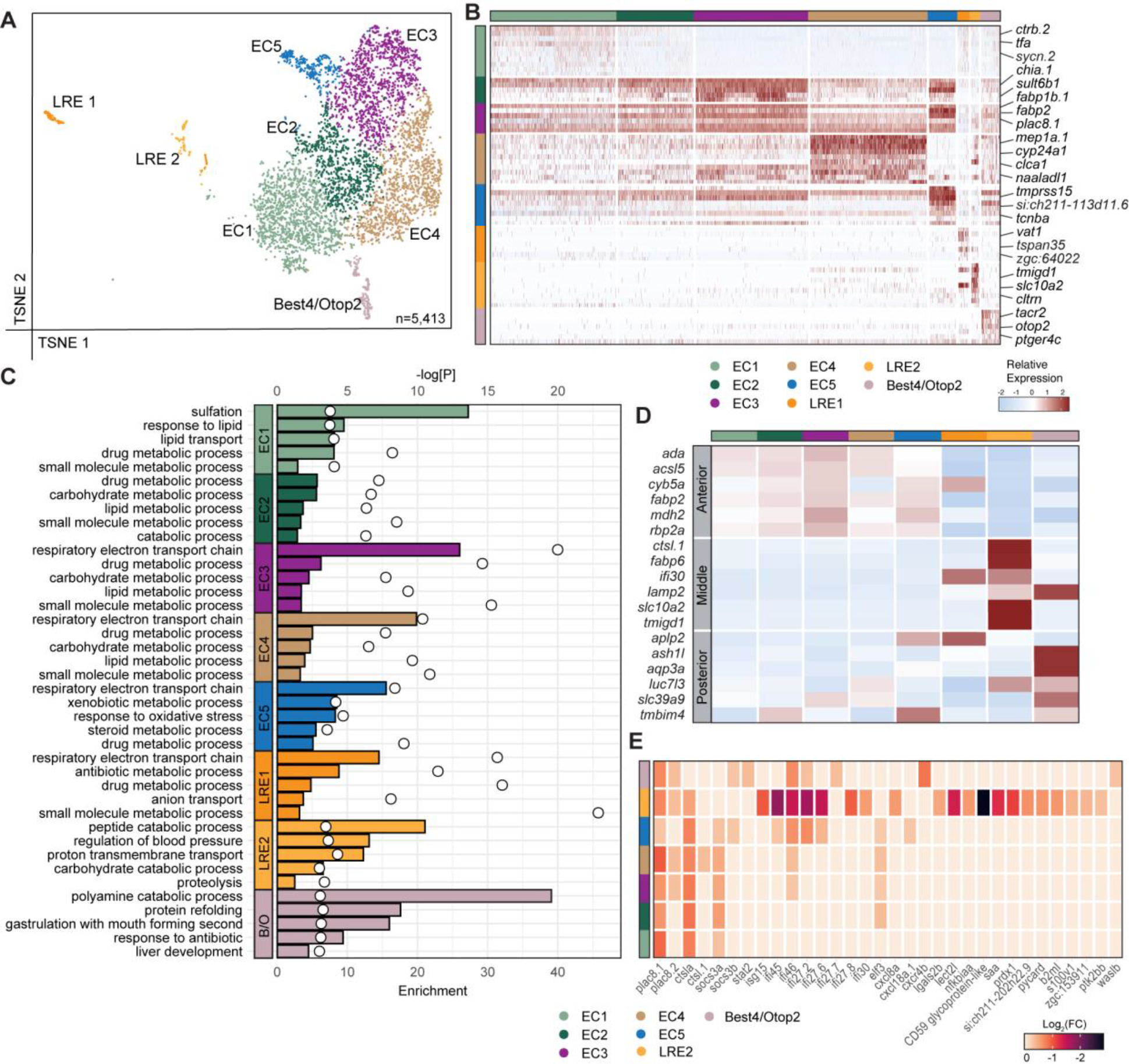
Microbes promote distinct immune signals in regionally specified absorptive cells. (*A*) t-SNE plot of absorptive cells color coded by cell state. (*B*) Heatmap of cluster markers for absorptive cells, colored by relative gene expression. Cell types are indicated by colored bars on the left and top. Several top markers for each cluster relative to other absorptive cell populations are shown on the right axis of the heatmap. (*C*) Gene ontology enrichment analysis of absorptive cells based on genetic markers from the conventional single-cell RNA sequencing dataset. Top 5 non-redundant GO terms are shown for each cell state. Enrichment score is represented by bar length and p-value is indicated with white circles. (*D*) Heatmap showing relative expression of established regional marker genes in each absorptive cell type. (*E*) Heatmap of differentially expressed immune related genes in GF relative to CV absorptive cells, color coded according to Log2(FC). All non-zero value expression changes are significant (p<0.05) as determined with a non-parametric Wilcoxon rank sum test. LRE 1 showed no significant differential immune gene expression and was therefore not included.

As absorptive cells make frequent, direct contacts with lumenal microbes, we expected substantial transcriptional shifts in response to microbe-free development. To test this hypothesis, we assessed the consequences of microbiome exposure on immune responses within absorptive populations. Enterocyte clusters one to four had uniform transcriptional responses to GF growth, including downregulation of *plac8.1*, *cathepsin La* (*ctsla*), and the interferon pathway element *socs3a* (Fig. 4E). By contrast, cluster five cells, a putative foregut IEC population, showed a robust response to GF growth that included suppressed induction of interferon-response genes, and diminished expression of the *cxcl18a.1* chemokine (Fig. 4E). LRE1 cells did not display significant changes in GF fish, consistent with renal localization. However, mid-intestinal LRE2 cells exhibited dramatic changes following microbial elimination, including suppressed expression of the NF-κB pathway elements *lgals2b* and *nfkbiaa* (Fig. 4E). Remarkably, GF LRE2 cells exhibited considerable similarities with the GF response of cluster one goblet cells, including significant downregulation of *prdx1*, *lect2l*, *saa*, and numerous interferon-stimulated genes (Fig. 4E), suggesting that mid-intestinal LRE2 and goblet one clusters have overlapping immune responses to microbial encounters. Collectively, our data indicate that mature zebrafish IECs include a sophisticated organization of absorptive lineages that contribute to regionally-specialized immune responses to gut microbes.

### Cell-specific effects of microbial exposure on leukocyte and stromal activity

As our data included non-epithelial lineages, we expanded our study to map relationships between microbiome colonization and gene activity in leukocytes and stromal cells, critical regulators of host-microbe interactions. We uncovered two highly distinct larval leukocyte clusters (Fig. 5A and Supplementary Table 1). Cluster one was a mixed phagocyte population that expressed macrophage and neutrophil markers such as *spi1b*, *mpeg1.1*, and *mpx* (Fig. 5B, C). In contrast, cluster two cells expressed classical markers of developing T cells (Ma et al., 2013), such as *ikaros* (*ikzf1*), *runx3* and *ccr9a*, as well as the *T-cell receptor alpha/delta variable 14.0* gene segment (*tradv14.0*) (Fig. 5B-D), supporting prior reports of immature T cells in 6 dpf fish (Ma et al., 2013). Future work is required to determine if T cells have already seeded the larval intestine by day six, or if these cells originated from thymic or kidney tissue attached to dissected guts. The microbiome primarily impacted gene expression within phagocytes, where GF growth led to significantly diminished expression of interferon and cytoskeletal components relative to CV controls (Fig. 5G), and attenuated production of key immune regulators such as *stat1a* and *stat2*, and the pro-inflammatory cytokines *il1b*, *tnfa* and *tnfb* (Fig. 5I).

**Figure 5.**
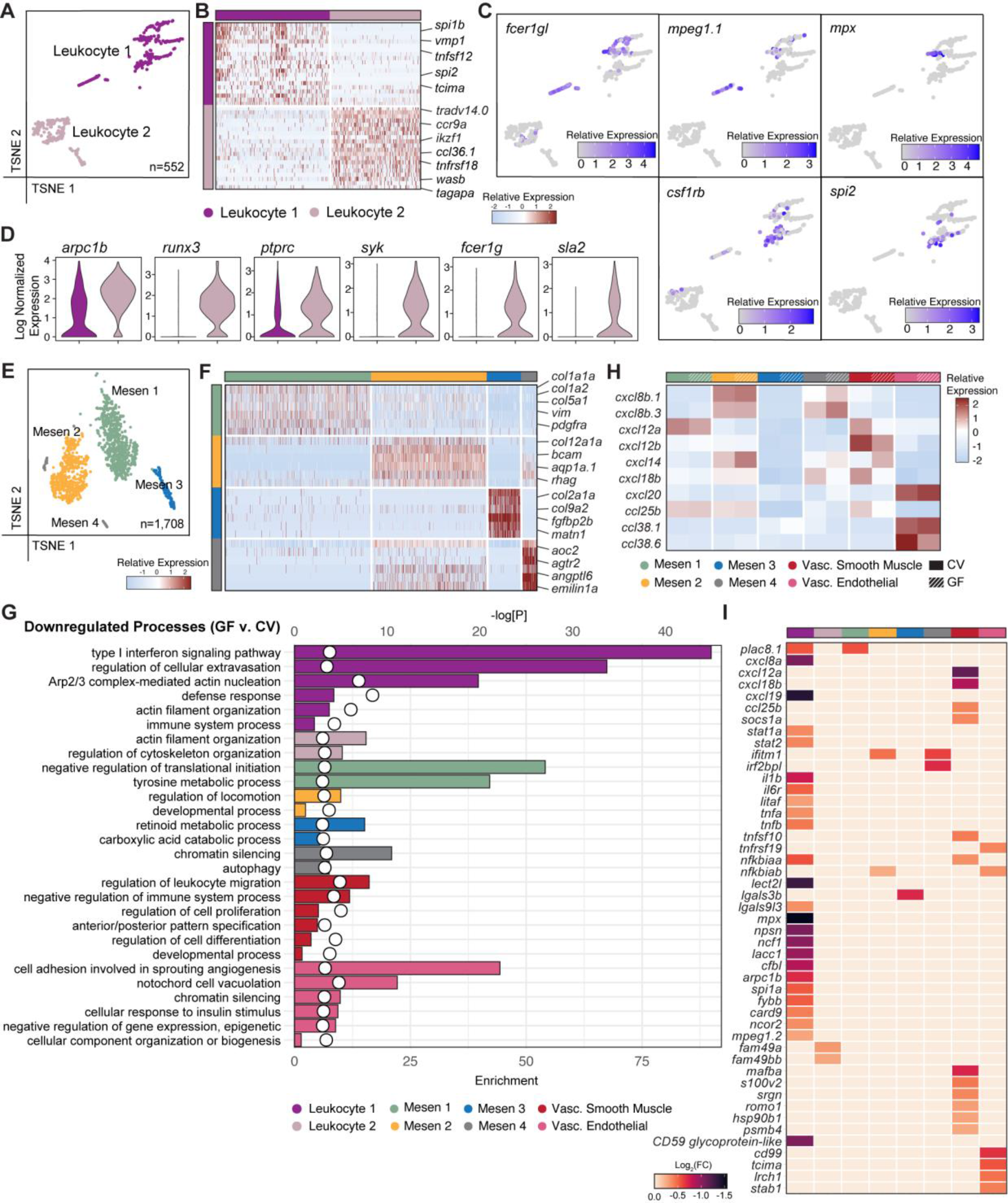
Stromal and leukocyte populations have subtype-specific responses to commensal microbes. (*A*) t-SNE plot of leukocytes, color coded by cell cluster. (*B*) Heatmap of cluster markers for leukocytes, colored by relative gene expression. Cell types are indicated by colored bars on the left and top. Several top cluster markers relative to the other leukocyte population are shown on the right axis of the heatmap. (*C*) t-SNE plots of leukocytes showing cell-specific expression of leukocyte subset markers. (*D*) Violin plots showing log normalized expression of marker genes for leukocyte 2 cells. (*E*) t-SNE plot of color-coded mesenchymal clusters. (*F*) Heatmap of cluster markers for mesenchymal cells, colored by relative gene expression. Cell types are indicated by colored bars on the left and top. Several top cluster markers relative to the other mesenchymal population are shown on the right axis of the heatmap. (*G*) GO enrichment analysis of downregulated genes in GF relative to CV stromal and leukocyte populations. Enrichment score is represented by bar length and p-value is indicated with white circles. (*H*) Heatmap showing relative expression of chemokines in CV or GF stromal and leukocyte subsets. (*I*) Heatmap of differentially expressed immune related genes in GF relative to CV leukocyte and stromal cell populations, color coded according to Log2(FC). All non-zero value expression changes are significant (p<0.05) as determined with a non-parametric Wilcoxon rank sum test.

Mesenchymal cells segregated into four distinct clusters (Fig. 5E), of which cluster one represented a fibroblast population that expressed extracellular matrix components such as *col1a1a* and *col1a2*, and the fibroblast marker *vimentin* (*vim*) (Fig. 5F). The identity of mesenchymal cluster two is unclear; however, it was marked by expression of ammonia transporter *rhag,* and *aqp1a.1* (Fig. 5F) involved in ammonia, water, and CO2 transport (Horng et al., 2015; Talbot et al., 2015), suggesting these cells regulate gas and ion movement. Cluster three cells were marked by ECM components *matrilin 1* (*matn1*) and several collagens, as well as *fibroblast growth factor binding protein 2b* (*fgfbp2b*), also indicative of fibroblast identity (Fig. 5F). Finally, mesenchyme cluster four was marked by expression of soluble pattern recognition receptors from the collectin family (Supplementary Table 1), and vasculature markers *angptl6* and *agtr2* (Fig. 5F), indicating that cluster four likely represents perivascular fibroblasts. Among the mesenchymal clusters, removal of the microbiome primarily attenuated expression of genes associated with metabolism (Fig. 5G). In contrast, GF growth had sizable effects on gene expression in vascular smooth muscle and endothelial cells. Relative to CV controls, GF vascular cells expressed significantly lower amounts of genes that regulate leukocyte migration, cell proliferation, and sprouting angiogenesis. (Fig. 5G). Furthermore, unlike mesenchymal cell-types, vascular smooth muscle cells exhibited significantly decreased chemokine expression under GF growth conditions (Fig. 5H-I), implicating vascular cells as an intermediary in microbe-dependent leukocyte recruitment. Consistent with a role for vascular cells in mediating microbial recruitment of leukocytes, we found that, compared to CV controls, vascular smooth muscle cells from GF fish downregulated expression of the lymphocyte chemotactic regulator *cxcl12b,* and the granulocyte chemotaxis regulator *cxcl18b*, while vascular endothelial cells downregulated *cd99*, a promoter of trans-endothelial leukocyte migration (Schenkel et al., 2002). In summary, we have identified distinct leukocyte and stromal cell subtypes in the larval gut. Our data uncover differential degrees of microbial sensitivity within the subtypes and implicate vascular cells as agents of microbe-responsive leukocyte migration.

### The microbiome is essential for intestinal vascularization

Integrated analysis of CV and GF data revealed microbe-dependent gene expression changes in vascular endothelial and smooth muscle populations, including significantly diminished expression of vasculature developmental regulators (Fig 5G-I). Thus, we reasoned that, like mice (Reinhardt et al., 2012; Stappenbeck et al., 2002), microbes may promote zebrafish intestinal angiogenesis. A closer look at vascular cells showed that larvae raised in GF conditions expressed lower amounts of pro-angiogenic factors such as *moesin a* (*msna*) and *cdh5*, as well as BMP regulators involved in vascular morphogenesis (He and Chen, 2005; Mouillesseaux et al., 2016), such as *smad5* and *smad6a* (Fig. 6A). Likewise, we detected significant drops in expression of *angptl4* and transcriptional regulators *pre-B-cell leukemia homeobox 1a* (*pbx1a*) and *pbx4* in vascular smooth muscle (Fig. 5A), known regulators of vascular development (Cvejic et al., 2011; Kao et al., 2015). Combined, these data raise the possibility that GF growth has detrimental consequences for formation of gut-associated vascular tissue. Intestinal vasculogenesis commences approximately three days after fertilization (Isogai et al., 2001), a time that matches microbial colonization of the lumen. At this stage, angioblasts migrates ventrally from the posterior cardinal vein, establishing the supra-intestinal artery, and a vascular plexus that gradually resolves into a parallel series of vertical vessels and the sub-intestinal vein(Goi and Childs, 2016; Lenard et al., 2015; Nicenboim et al., 2015). The gut vasculature delivers nutrients from the intestine to the hepatic portal vein, supporting growth and development. To determine if the microbiome affects intestinal vasculogenesis, we used *kdrl:mCherry* larvae to visualize the vasculature of fish that we raised in the presence, or absence of a conventional microbiome for six days (Fig. 6B-C). We did not observe effects of the microbiome on formation or spacing of the supra-intestinal artery and the sub-intestinal vein, VEGF-independent processes. In both groups, the artery and vein effectively delineated the dorsal and ventral margins of the intestine (Fig. 6B and 6C). In contrast, removal of the microbiome had significant effects on development of connecting vessels, a VEGF-dependent event. Consistent with this, we observed a near 50% reduction of intestinal *kdrl:mCherry* signal in GF larvae compared to CV counterparts (Fig. 5D). Thus, we conclude that microbial factors are essential for proper development of the zebrafish intestinal vasculature.

**Figure 6.**
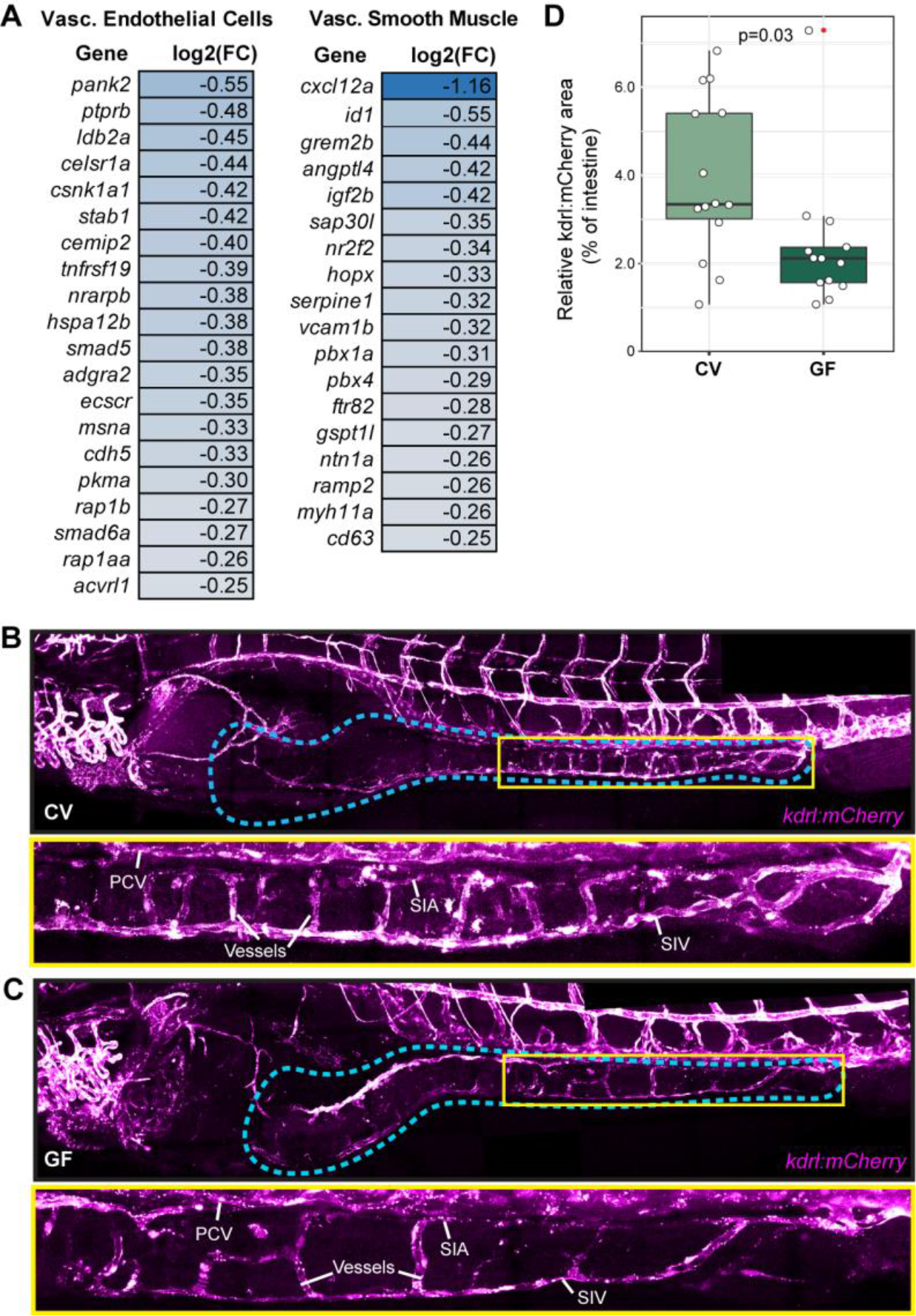
Microbes promote pro-angiogenic factor expression and intestinal vasculogenesis. (*A*) Downregulated expression of pro-angiogenic factors in GF relative to CV vascular endothelial and vascular smooth muscle populations (p<0.05). Significance was determined with a non-parametric Wilcoxon rank sum test. (*B-C*) Expression of *kdrl:mCherry* in zebrafish 6 dpf raised under CV (B) or GF (C) conditions. Corresponding brightfield images were used to identify the intestine, outlined in blue. Bottom panels in B and C show enlarged region of middle to posterior intestine within yellow boxes of respective upper panels. PCV- posterior cardinal vein; SIA- supra-intestinal artery; SIV- sub-intestinal vein. (*D*) Box and whisker plot showing the area of intestinal kdrl:mCherry signal relative to total intestinal area. n=14 and n=13 for CV and GF fish respectively. Outlier is indicated with red dot. Significance was determined via Student’s *t* test.

## DISCUSSION

Gut microbial factors are critical determinants of animal development (Sekirov et al., 2010). Comparative studies with CV and GF zebrafish larvae uncovered numerous microbial effects on the host, including impacts on proliferation, cell fate specification, and metabolism (Bates et al., 2006; Hooper et al., 2001; Rawls et al., 2004; Reikvam et al., 2011). Importantly, the molecular and genetic networks that determine intestinal development are highly similar between zebrafish and mammals (Davison et al., 2017; Heppert et al., 2021; Lickwar et al., 2017). Thus, discoveries made with fish have the potential to reveal foundational aspects of host-microbe relationships. However, important knowledge gaps prevent us from maximizing the value of zebrafish-microbe interaction data. In particular, we know less about cellular composition within the zebrafish intestinal epithelium compared to mice and humans. For example, the zebrafish epithelium contains basal cycling cells that likely act as epithelial progenitors (Crosnier et al., 2005; Li et al., 2020; Peron et al., 2020; Rawls et al., 2004; Wallace et al., 2005). However, we lack genetic markers that allow us to identify or isolate the progenitor population for experimental characterization. Similarly, the extent of functional heterogeneity with absorptive epithelial lineages requires clarification. To bridge these deficits, and permit cell-by-cell definition of host responses to the microbiome, we prepared single-cell atlases of the larval intestine raised under conventional or germ-free conditions. We identified thirty-five transcriptionally distinct clusters in the gut and associated tissue, including cells that have not been described to date. Comparisons between conventional and germ-free fish allowed us to delineate impacts of the microbiome on growth, patterning, immune, and metabolic processes in each cell type. We believe these findings constitute a valuable resource that will support efforts to understand how microbes influence vertebrate physiology. To make our data publicly accessible for single-cell gene expression analysis under conventional and germ-free conditions, we have uploaded our datasets to the Broad Single Cell Portal, a web-based resource for single cell visualization Looking at the intestinal epithelium, we identified cycling cells that express classical intestinal stem cell markers, such as the Delta-like ligand *dld*, the Notch pathway components *ascl1a* and *atoh1b,* and Notch-responsive *hes-related* transcription factor family members, such as *her2* and *her15.1*. Notably, expression of several cell cycle markers, *her2*, *her15.1*, and *dld* declined in this population in GF larvae, consistent with diminished epithelial growth in fish raised in the absence of a microbiome (Bates et al., 2006; Rawls et al., 2004). Our data align with related work in vertebrates and invertebrates (Buchon et al., 2009; Cheesman et al., 2011; Rawls et al., 2004; Reikvam et al., 2011), and raise the possibility that we may have uncovered intestinal stem cell markers for zebrafish larvae. In mammals, Lgr5 is a classical intestinal stem cell marker, with roles in Wnt signaling (Haegebarth and Clevers, 2009). Though Wnt is important for microbe-dependent intestinal epithelial growth in zebrafish (Cheesman et al., 2011), the zebrafish genome does not appear to encode an *lgr5* ortholog. Instead, the fish genome encodes related *lgr4* and *lgr6* genes (Hirose et al., 2011). We did not observe expression of *lgr6* in the fish gut and detected *lgr4* expression primarily in absorptive cells, raising the possibility that Wnt-Lgr signaling may not be essential for specification and growth of fish intestinal stem cells. In this regard, zebrafish may be more akin to *Drosophila*, where Wnt activity has ancillary roles in midgut intestinal stem cell growth (Cordero et al., 2012; Lee et al., 2009; Lin et al., 2008; Tian et al., 2016). In the future, it will be of interest to perform lineage tracing with candidate progenitor cells identified in this study to test their ability to generate a mature epithelium, and to resolve the roles of Wnt-Lgr activity in zebrafish epithelial homeostasis.

Separate to cycling, Notch-positive cells, we identified transcriptional markers for secretory enteroendocrine cells, and mucin-producing goblet cells (Crosnier et al., 2005; Ng et al., 2005; Wallace et al., 2005). Examination of the enteroendocrine population revealed a sophisticated arrangement of lineages that can be distinguished based on their rostro-caudal distribution and on the profile of peptide hormones they produce, indicating a spatially complex pattern of hormone production within the developing intestinal epithelium. Integrated comparisons between CV and GF fish uncovered pronounced effects of the microbiome on goblet cell immune signaling. We found that gut microbes induce goblet-specific expression of the interferon-responsive transcription factor *stat2*, several interferon-inducible genes, and genes involved in leukocyte chemotaxis (*lect2l*, *saa*, *ccl19a.1*, *ccl19b*), implicating goblet cells in the coordination of host immune responses to a conventional microbiome.

Upon examination of differentiated lineages, we identified transcriptional signatures of regionally and spatially specialized enterocytes in CV larvae, as well as an extensive profile of gene expression in lysosome-rich enterocytes that mediate protein absorption and metabolism (Park et al., 2019). Our findings provide a molecular underpinning of the regionally controlled nature of nutrient metabolism in the zebrafish gut. Perhaps more intriguingly, we also uncovered two lineages that were unknown in zebrafish. Specifically, we identified Best4/Otop2-positive enterocytes that are likely enriched in the posterior intestinal epithelium. Best4/Otop2 cells are a minimally characterized cell type only recently discovered in the human colon (Parikh et al., 2019; Smillie et al., 2019). Given the utility of zebrafish for examination of gut development, particularly in the context of host-microbe interactions, we believe fish will be of considerable value for *in vivo* characterization of Best4/Otop2 cells. Separately, we identified cells that are enriched for expression of markers highly associated with intestinal tuft cells, including the *pou2f3* master regulator. Tuft cells are a relatively under-characterized cell type that share developmental trajectories with secretory lineages, but appear to activate mucosal type II immune responses. At present, it is unclear if zebrafish tuft-like cells are involved in mucosal defenses. Nonetheless, our identification of Best4/Otop2 and tuft-like cells within the zebrafish intestinal epithelium underscores the similarities between fish and mammalian intestines. We further note that our datasets are in broad agreement with a recent study reporting single-cell profiles for sorted zebrafish IECs (Wen et al., 2021).

The availability of a high-resolution transcriptional atlas of the zebrafish intestine allowed us to map microbial effects on each cell type. While it is possible that some of the cell-specific changes observed result from GF derivation, we find that our recapitulation of known microbe-responsive processes makes this unlikely to be a major confounding factor. Importantly, the resolution provided by single-cell approaches allowed us to uncover a large number of unknown microbe-driven processes in the host, and resolve each process to the level of distinct cell clusters. Our work shows that microbiota-dependent control of growth, developmental, metabolic and immune processes display remarkable cellular specificity. To provide one example, we will discuss effects of the microbiota on host immune activity; however, we note our data permit identification of microbial impacts on many physiological processes.

Our work revealed a hitherto unknown complexity of germline-encoded immune gene expression patterns in CV fish, suggesting a refined partitioning of immune functions among intestinal epithelial cell types. Absorptive intestinal epithelial cells expressed enriched amounts of detoxifying alkaline phosphatases (Bates et al., 2006), and myeloid-activating *serum amyloid A* (Kanther et al., 2011; Murdoch et al., 2019). In contrast, progenitor and tuft cells expressed elevated levels of the bacterial peptidoglycan sensor *nod2* and core NF-kB pathway elements, whereas enteroendocrine cells and leukocytes expressed larger amounts of *il22*, a cytokine that activates epithelial defenses (Dudakov et al., 2015). Phagocytes were characterized by elevated expression of pro-inflammatory cytokines such as *il1b, tnfa* and *tnfb*, whereas mesenchymal cells were prominent sources of immune-regulatory TGF-beta class cytokines. Comparisons between CV and GF fish uncovered a remarkable input from the microbiome on all these processes, with cell-specific expression of many immune effectors and mediators declining, relocating, or disappearing almost entirely in GF fish. Future work will be needed to elucidate impacts of cell-specific immune signals on intestinal homeostasis.

To test developmental consequences of microbial removal on larvae, we focused on intestinal vasculogenesis. In fish, the intestinal vasculature arises from angioblasts that migrate ventrally from the posterior cardinal vein, and establish a plexus that gradually resolves into the dorsal supra-intestinal artery, the ventral sub-intestinal vein, and a series of parallel vessels that connect artery and vein (Goi and Childs, 2016; Isogai et al., 2001; Lenard et al., 2015; Nicenboim et al., 2015). We noted diminished expression of key angiogenesis regulators in GF larvae, particularly VEGF-class receptors with established roles in formation of connecting vessels (Goi and Childs, 2016). Examination of GF fish showed that the microbiota is dispensable for positioning and spacing of the artery and vein. In contrast, removal of the microbiota had deleterious effects on connecting vessels, confirming a role for the microbiome in establishing the intestinal vasculature. Our results match observations from mice, where germ-free growth also diminishes villus angiogenesis (Reinhardt et al., 2012; Stappenbeck et al., 2002), suggesting a shared requirement for microbial cues to direct intestinal angiogenesis in vertebrates. We believe the advances made in this study will allow us to trace the molecular, and cellular networks that control intestinal vasculogenesis in a developing vertebrate.

## ACKNOWLEDGEMENTS

We acknowledge flow cytometry support from Dr. Aja Rieger and Sabina Baghirova, as well as support with single-cell library preparation from Dr. Joaquin Lopez-Orozco. Flow Cytometry Facility Experiments were performed at the University of Alberta Faculty of Medicine & Dentistry Flow Cytometry Facility, which receives financial support from the Faculty of Medicine & Dentistry and Canada Foundation for Innovation (CFI) awards to contributing investigators. We also acknowledge imaging help from Dr. Xuejun Sun of the Department of Oncology Cell Imaging Facility and Arlene Oatway of the Department of Biological Sciences Microscopy Facility at the University of Alberta. We would also like to thank Science facility. This work was supported by grants from the Canadian Institute of Health Research (Grant # PJT 159604). RJW has funding support through the University of Alberta Faculty of Graduate Studies and Research, National Science and Engineering Research Council Graduate Scholarships, and Alberta Innovates Graduate Student Scholarships.

## MATERIALS AND METHODS

### Data Availability

Cell Ranger raw output files are available from the NCBI GEO database (GSE161855). Processed data is available for visualization and analysis on the Broad Single Cell Portal (SCP1623).

### Zebrafish strains and maintenance

Zebrafish were raised and maintained using protocols approved by the Animal Care & Use Committee: Biosciences at the University of Alberta, operating under the guidelines of the Canadian Council of Animal Care. TL strain zebrafish were used for single-cell RNA sequencing, Nanostring gene expression analysis, and transmission electron microscopy, and the *Tg*(*kdrl:mCherry*) line (Wang et al., 2010a) was used for analysis of intestinal vasculogenesis. Adult fish were raised and maintained within the University of Alberta fish facility at 28°C under a 14 hour/ 10 hour light/ dark cycle as previously described (Westerfield, 2000). For larval analysis, breeding tanks were set up overnight with 1 male and 1 female separated by a divider until morning. Fish were bred for 1 hour, then embryos were collected, rinsed gently with facility water, and transferred to culture flasks (Corning) with 15 mL embryo media (EM) (prepared as in Melancon et al., 2017) and 15 embryos per flask. Embryos were raised at 29°C under a 14 hour/ 10 hour light/ dark cycle until 6 days post fertilization.

### Generating germ-free zebrafish

Fish embryos were made germ-free essentially as in (Melancon et al., 2017). A clutch of embryos was collected then washed and split into two cohorts. The CV cohort was kept in sterile EM, while the GF cohort was kept in sterile EM supplemented with ampicillin (100 µg/mL), kanamycin (5 µg/mL), amphotericin B (250 ng/mL), and gentamicin (50 µg/mL). Embryos were washed every 2 hours with EM or EM plus antibiotics for CV and GF cohorts respectively. Once at 50% epiboly, the GF cohort was successively washed three times in EM, then 2 minutes in 0.1% polyvinylpyrrolidone-iodine (PVP-I) in EM, followed by three EM washes, then a 20-minute incubation with 0.003% sodium hypochlorite (bleach) in EM. Embryos were washed three more times then transferred into tissue culture flasks with sterile EM. The CV cohort received the same number and duration of washes, using EM in lieu of dilute PVP-I or bleach. All work was performed in a biosafety cabinet sterilized first with 10% bleach, followed by 70% ethanol. We tested for bacterial contamination in GF flasks at 4 days post-fertilization, according to established protocol (Melancon et al., 2017). EM was collected from CV and GF culture flasks to test for bacteria by plating on TSA, as well as PCR against bacterial 16S rDNA. Parental tank water and sterile filtered water were used as a positive and negative control respectively, where bacteria were positively identified in parental tank water and confirmed absent from sterile water. CV and GF flasks with bacteria present or absent respectively were used for subsequent analysis.

### Imaging and quantifying intestinal vasculature

*Tg(kdrl:mCherry)* fish (Wang et al., 2010a) were raised under CV or GF conditions for 6 dpf, then euthanized with tricaine and fixed overnight at 4 °C. Larvae were washed 3X in PBS then embedded in 0.7% UltraPure low melting point agarose (Invitrogen 16520) on a glass bottom dish. Tile and Z-stack images (5 µm sections) of whole fish were captured on a Leica Falcon SP8 equipped with a 25x 0.95NA Water HC Fluotar objective lens. Images were stitched with Leica Application Suite X software (Leica) and imported to FIJI to produce maximum intensity Z-projection images that were adjusted for brightness and contrast, as well as false color manipulations. To quantify intestinal vasculature, corresponding brightfield images were used to set intestinal boundaries in FIJI. Fluorescent images were then converted to binary images and the area of kdrl:mCherry signal relative to the area of the whole intestine was measured. Box plots were generated in RStudio with ggplot2.

### Generation and analysis of transient transgenic zebrafish

*Tg(tnfrsf11a:GFP)* zebrafish were generated using the Tol2kit (Kwan et al., 2007). Briefly, a 3441 base pair fragment upstream of the of the *tnfrsf11a* transcription start site was amplified by PCR from zebrafish genomic DNA, then subcloned into the 5’ entry vector using KpnI and SacII restriction sites. The p5E-3.4*- tnfrsf11a* construct was confirmed via restriction digest, and gateway cloning was used to combine the 5’ entry, middle entry (pME-EGFP), and 3’ (p3E-polyA) entry clones into the destination vector (pDestTol2CG2). The final construct was confirmed via restriction digest. To generate transient transgenics, 1-cell stage embryos were injected with approximately 50 pg DNA and 25 pg transposase RNA. Injected embryos were raised to 6 dpf, and larvae were screened for both *cmlc2:GFP* expression and intestinal GFP signal. Positive larvae were euthanized in 5X tricaine and intestines were dissected into ice-cold 4% PFA in PBS and fixed overnight at 4°C. Guts were then washed in PBS + 0.75% Triton-X (PBT), blocked in PBT + 3% BSA (PBTB) for 1 hour, then incubated in 1° antisera (Invitrogen chicken anti-GFP, 1:4000) in blocking solution overnight at 4°C. The next day, guts were washed in PBT, incubated with 2° antibody (1:1500) and Alexa Fluor^TM^ 647 Phalloidin (Invitrogen A22287, 1:2000) in PBTB for 1 hour. Guts were then washed and counterstained with Hoechst 33258 (ThermoFisher, H3569, 1:2000 dilution) in PBT before mounting. Z stack images (0.3-0.5 mm sections) were acquired using an Olympus IX-81 spinning disc confocal equipped with a Hamamatsu EMCCD (C9100-13) camera and operated with Volocity 4. Z stack images were exported and processed in Fiji (Schindelin et al., 2012).

### Generating single-cell suspensions for single cell RNA-seq

Fish from the same embryo clutch were derived CV or GF as described. Five larvae were euthanized at a time in PBS plus 5X tricaine, then intestines were immediately dissected with sterilized equipment and placed into 200 µL sterile PBS in a 1.5 mL microfuge tube on ice, alternating five CV and five GF intestines until 25 intestines (replicate 1) or 55 intestines (replicate 2) were collected per condition (80 intestines total per condition). Total dissection time was kept below 2 hours for each replicate. Immediately following dissections, intestines were incubated in 1.5 mL microfuge tubes with 200 µL of dissociation cocktail containing 1 mg/mL fresh collagenase A, 40 µg/mL proteinase k, and 0.25% trypsin in PBS for 30 minutes at 37°C, pipetting up and down 40X every 10 minutes to aid digestion. Then, either (Replicate 1) ZombieAqua viability dye (BioLegend) was added at the beginning of dissociation to a final concentration of 1:1000 to stain dead and dying cells, 10% non-acetylated BSA in PBS was added to the dissociation cocktail (final concentration of 1%) to stop digestion, cells were spun for 15 minutes at 0.3 RCF and 4 °C to pellet cells, cells were gently re-suspended in 200 µL PBS+0.04% BSA (non-acetylated) and spun down through a 40 µm cell strainer (Pluriselect) at 0.3 RCF for 1 minute at 4 °C, then filtered cells were sorted on a BD FACS Aria III to collect live single cells (ZombieAqua negative); or (Replicate 2) 10% non-acetylated BSA in PBS was added to the dissociation cocktail (final concentration of 1%) to stop digestion, and the cells were spun for 15 minutes at 0.3 RCF and 4 °C to pellet cells. Cells were then gently re-suspended in 200 µL PBS+0.04% non-acetylated BSA and spun down through a 40 µm cell strainer (Pluriselect) at 0.3 RCF for 1 minute at 4°C. Live cells were collected using OptiPrepTM Density Gradient Medium (SIGMA, D1556-250ML). Briefly, a 40% (w/v) iodixanol working solution was prepared with 2 volumes of OptiPrepTM and 1 volume of 0.04 %BSA in 1XPBS/DEPC-treated water. This working solution was used to prepare a 22% (w/v) iodixanol solution in the same buffer. One volume of working solution was mixed with 0.45 volume of cell suspension via gentle inversion. The solution mixture was transferred to a 15ml conical tube then topped up to 6 ml with working solution. The solution was overlaid with 3 ml of the 22% (w/v) iodixanol and the 22% iodixanol layer was overlaid with 0.5 ml of PBS+0.04% BSA. Viable cells were separated by density gradient created by centrifuging at 800xg for 20 min at 20°C. Viable cells were harvested from the top interface, which was then diluted in PBS+0.04% BSA. Live cells were pelleted at 0.3 RCF for 10 min at 4°C. Supernatant was decanted and cells were resuspended in PBS+0.04% BSA. (Both Replicates): Cell suspensions were then counted with a hemocytometer. Viability, as determined with Trypan blue, was >95% for all CV and GF samples. The single cell suspensions were immediately run through the 10X Genomics Chromium Controller with Chromium Single Cell 3′ Library & Gel Bead Kit v3.1. Libraries were constructed according to 10X Genomics Chromium Single Cell 3’ Library & Gel Bead Kit v3 protocol. Libraries were sent to Novogene, where QC was performed by Nanodrop for quantitation, agarose gel electrophoresis to test for library degradation/ contamination, and Agilent 2100 analysis for library integrity and quantitation. Paired-end sequencing was performed on the Illumina Hiseq platform with a read length of PE150 bp at each end.

### Processing and analysis of single cell RNA-seq data

For single cell analysis, Cell Ranger v3.0 (10X Genomics) was used to demultiplex raw base call files from Illumina sequencing and to align reads to the Zebrafish reference genome (Ensembl GRCz11.96). Cell Ranger output matrices were analyzed using the Seurat R package version 3.1.1 (Butler et al., 2018) in RStudio. Cells possessing fewer than 200 unique molecular identifiers (UMis), greater than 2500 UMIs, or greater than 50% mitochondrial reads were removed to reduce the number of low-quality cells and doublets. Seurat was then used to normalize expression values and perform cell clustering on integrated datasets at a resolution of 1.0 with 26 principal components (PCs), where optimal PCs were determined using JackStraw scores (Macosko et al., 2015) and elbow plots. After using the “FindMarkers” function in Seurat to identify marker genes for each cluster, clusters were annotated according to known cell type markers in zebrafish, or orthologous markers in mammals.

### Gene ontology (GO) enrichment analysis

Marker genes (p-value cut-off < 0.05), as well as down-regulated gene lists from the integrated dataset (p-value cut-off < 0.05) were analyzed in GOrilla (*Gene Ontology enRIchment anaLysis and visuaLizAtion tool)* to determine GO term enrichment (Eden et al., 2009). Genes were analyzed in a two-list unranked comparison using the whole dataset gene list as background. To remove redundant GO terms, enriched terms with associated p-values from GOrilla were run through REVIGO (REduce and VIsualize Gene Ontology) using SimRel semantic similarity metric with an allowed similarity of 0.4 (Supek et al., 2011). Bar plots were manually generated using ggplot2 in RStudio.

### NanoString nCounter gene expression analysis

Fish from the same embryo clutch were derived CV or GF as described. Fifteen 6 dpf zebrafish were taken per flask, with four replicates per condition. Larvae were euthanized in PBS plus 5X tricaine, then intestines were immediately dissected with sterile equipment and placed into 250 µL Trizol in a 1.5 mL microfuge tube on ice. Once 15 intestines were collected, samples were homogenized and stored at - 80°C. After freezing, samples were thawed, and standard Trizol-chloroform extraction was used to isolate RNA. Sample concentrations and quality were measured on an Agilent Bioanalyzer 2100 prior to shipping to NanoString Technologies for gene expression analysis using the nCounter® Elements^TM^ platform.

### 16S rRNA gene sequencing

Five days post fertilization, larval intestines were dissected, using aseptic technique, and collected in 200 ul of Microbead Solution. A total of thirty guts were collected, with ten guts pooled per replicate. The MoBio UltraClean Microbial DNA Isolation kit (Cat No. 12224-250) was used to extract microbial DNA. To assess the intestinal bacterial community composition, the V4 variable region of the 16s rRNA gene encompassed by the 515 forward primer and 806 reverse primer was sequenced. Quality control and sequencing was performed by Novogene Corporation using illumina Novaseq Platform PE250. Sequences were processed with QIIME2-2019.10 (qiime2.org). The DADA2 pipeline was used to join paired-end reads, remove chimeric sequences, and to generate the feature table used to resolve amplicon sequence variants using default parameters. DADA2 denoising resulted in 468,380 reads. Amplicon sequence variants (ASVs) represented by fewer than 200 reads across all samples were removed. A naïve Bayes classifier trained on SILVA132_99% full-length reference sequences was used to assign taxonomy. Taxonomy assignments were verified using NCBI blast. The sequence table was then filtered to exclude any sequences that were unassigned, not assigned past phylum level, or assigned as eukaryota, resulting in 441,712 sequences corresponding to 59 unique features.

### Transmission electron microscopy of adult zebrafish intestines

Adult fish were euthanized and dissected in accordance with protocols approved by the Animal Care & Use Committee: Biosciences at the University of Alberta, operating under the guidelines of the Canadian Council of Animal Care. To prepare samples for TEM, the posterior intestine was isolated and fixed in 2.5% glutaraldehyde, 2% PFA and 0.1M phosphate buffer solution for several days. Samples were then washed in 0.1M phosphate buffer, treated in 1% osmium tetroxide in 0.1M phosphate buffer, followed by additional washes. Intestines were subsequently dehydrated through a graded ethanol series, followed by infiltration with Spurr’s resin. Infiltrated samples were then embedded in flat molds in Spurr’s resin and cured overnight at 70°C. Blocks were then sectioned (70-90 nm thickness) on a Rechert-Jung UltracutE Ultramicrotome, and sections were stained with uranyl acetate, followed by lead citrate. Images were acquired using a FEI-Philips Morgagni 268 Transmission Electron Microscope operating at 80 kV and equipped with a Gatan Orius CCD camera.

## SUPPLEMENTARY MATERIAL

**Table S1.**
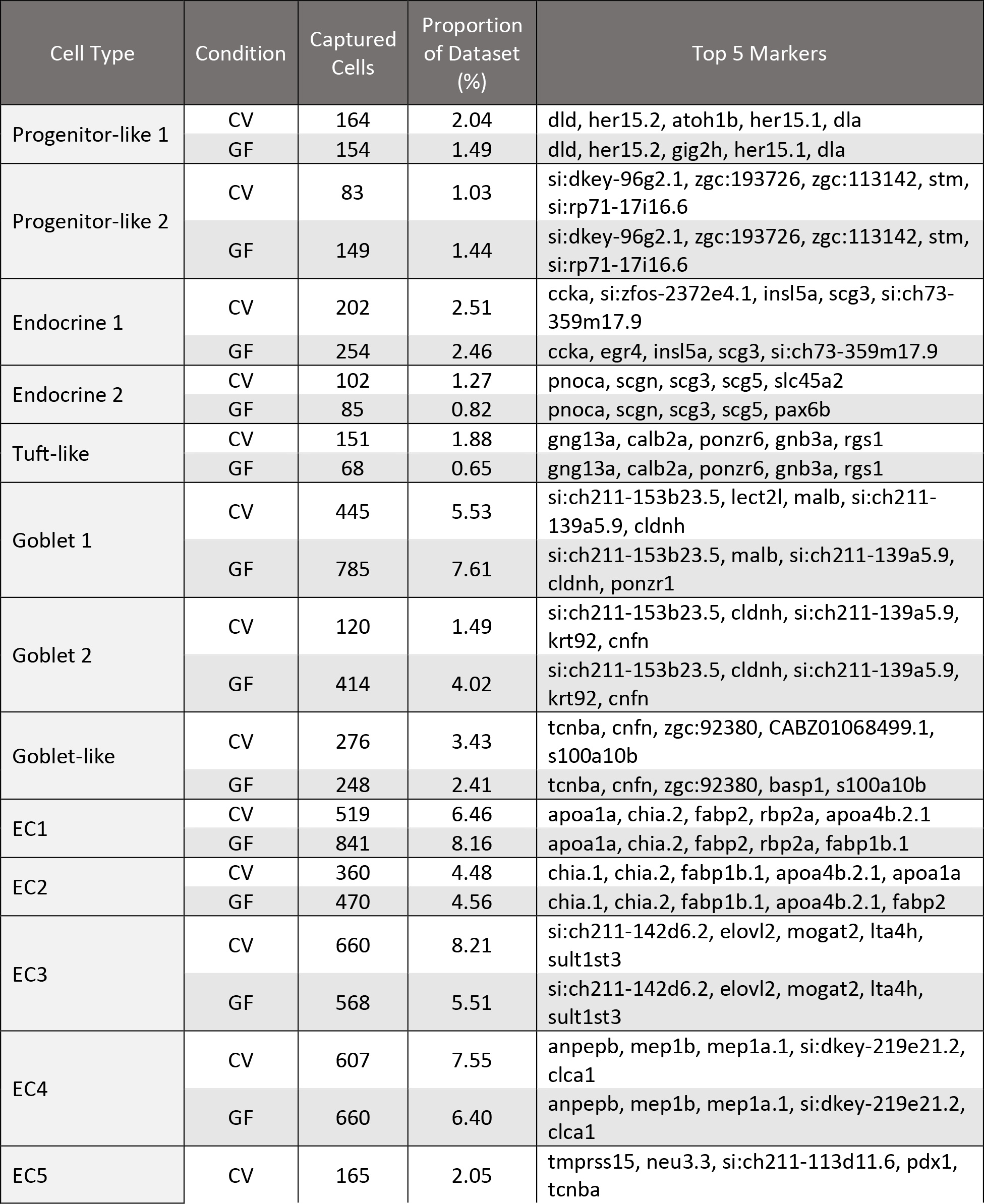
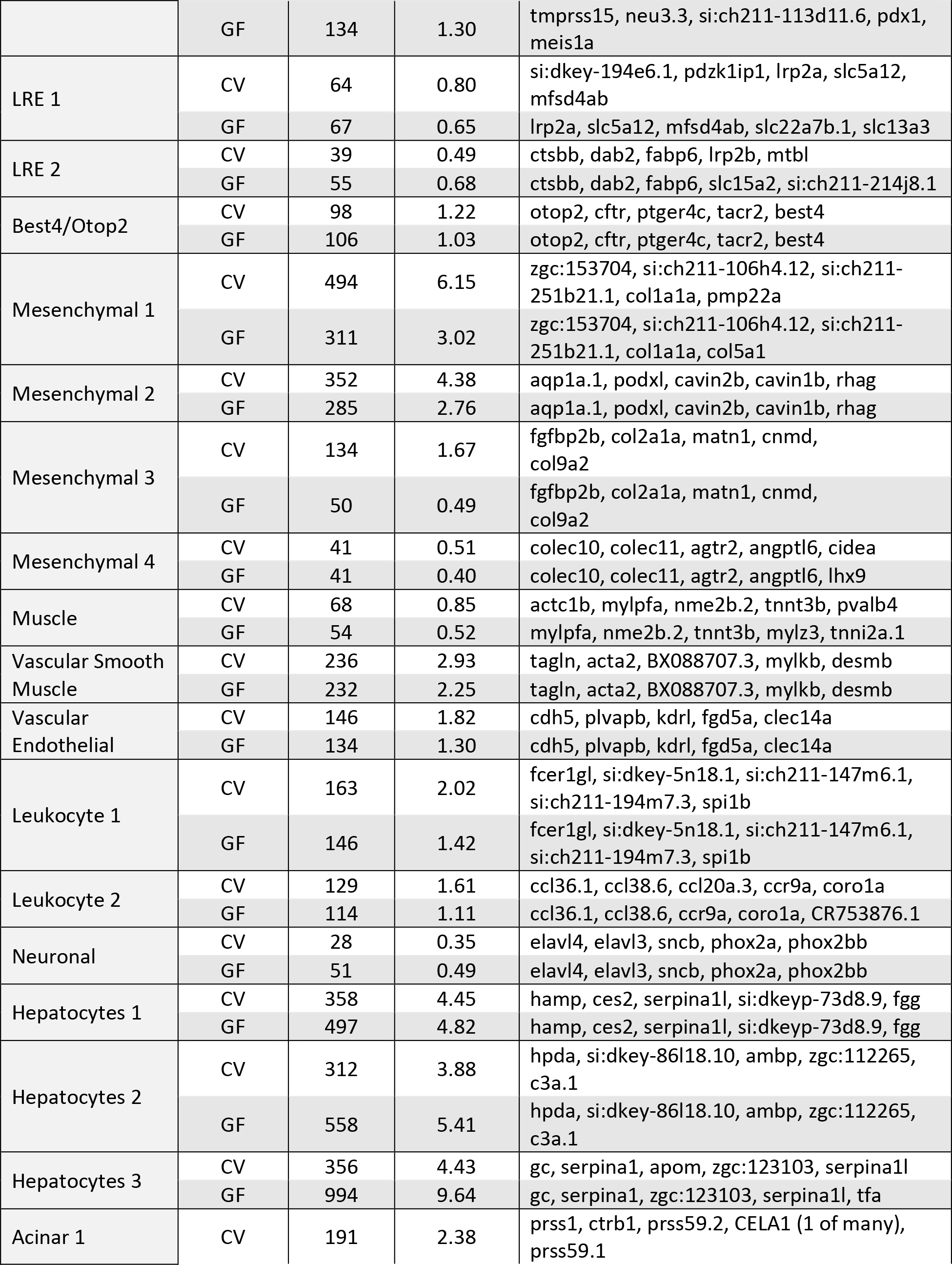
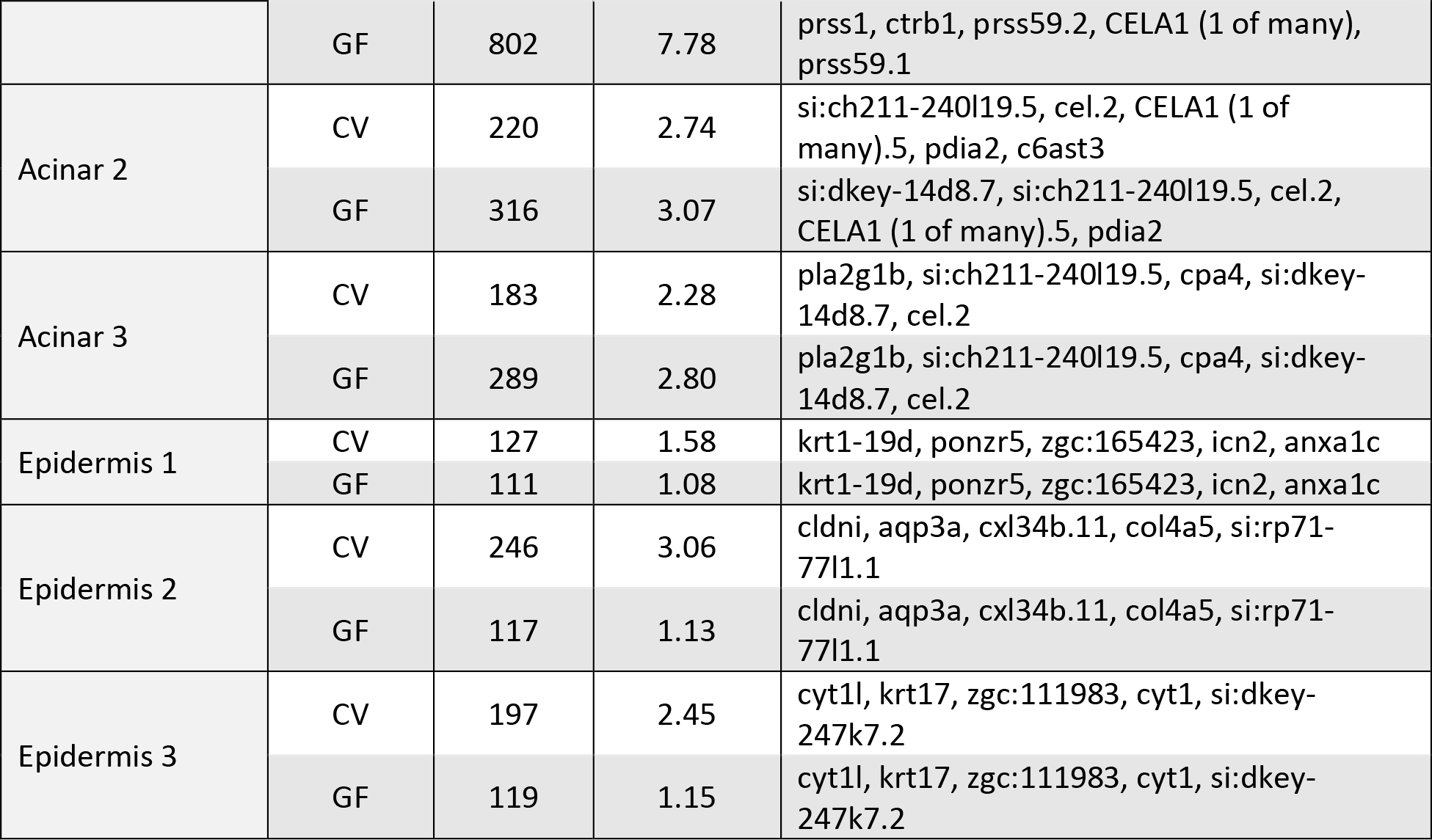
Single cell dataset composition and identifiers.

**Table S2.** Classification of 16S rRNA gene sequence datasets. Relative abundance is shown after removal of taxa that were <1% abundant.

| Phylum | Firmicutes |  | Proteobacteria |  |  |  |  |  |  |
| --- | --- | --- | --- | --- | --- | --- | --- | --- | --- |
| Class | Bacilli |  | ?-proteobacteria |  | ?-proteobacteria |  |  |  |  |
| Order | Bacillales | Lacto-bacillales | Aceto-bacterales | Rhizo-biales | Aeromo-nadales | Alteromo-nadales | Betaproteo-bacteriales | Pseudo-monadales | Vibrio-nales |
| CV1 | 0.78 | 6.20 | 4.91 | 0.25 | 0.72 | 0.46 | 0.84 | 1.64 | 84.21 |
| CV2 | 1.07 | 22.25 | 6.26 | 2.90 | 12.32 | 2.71 | 4.09 | 12.37 | 36.03 |
| CV3 | 0.56 | 35.06 | 2.76 | 1.18 | 3.75 | 2.28 | 4.67 | 6.06 | 43.69 |

**Table S3.**
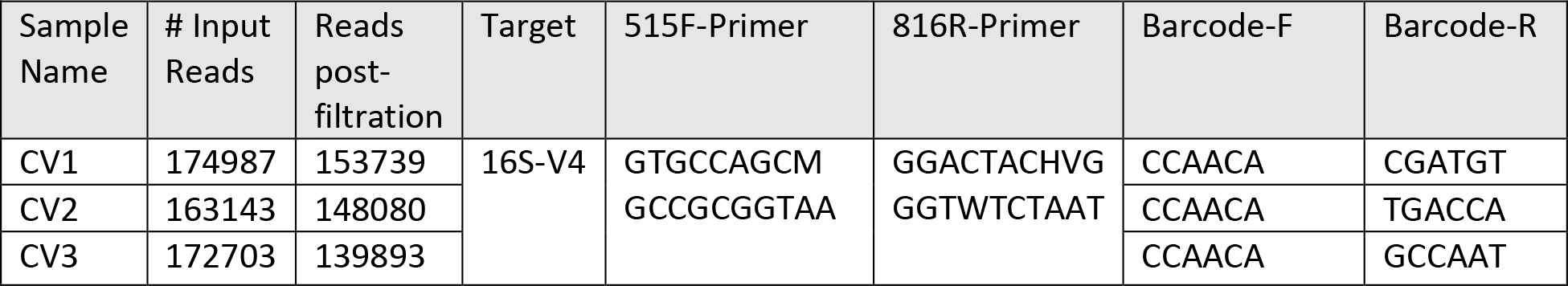
Description of 16s rRNA gene sequencing datasets used in this study and associated metadata.

**Table S4.**
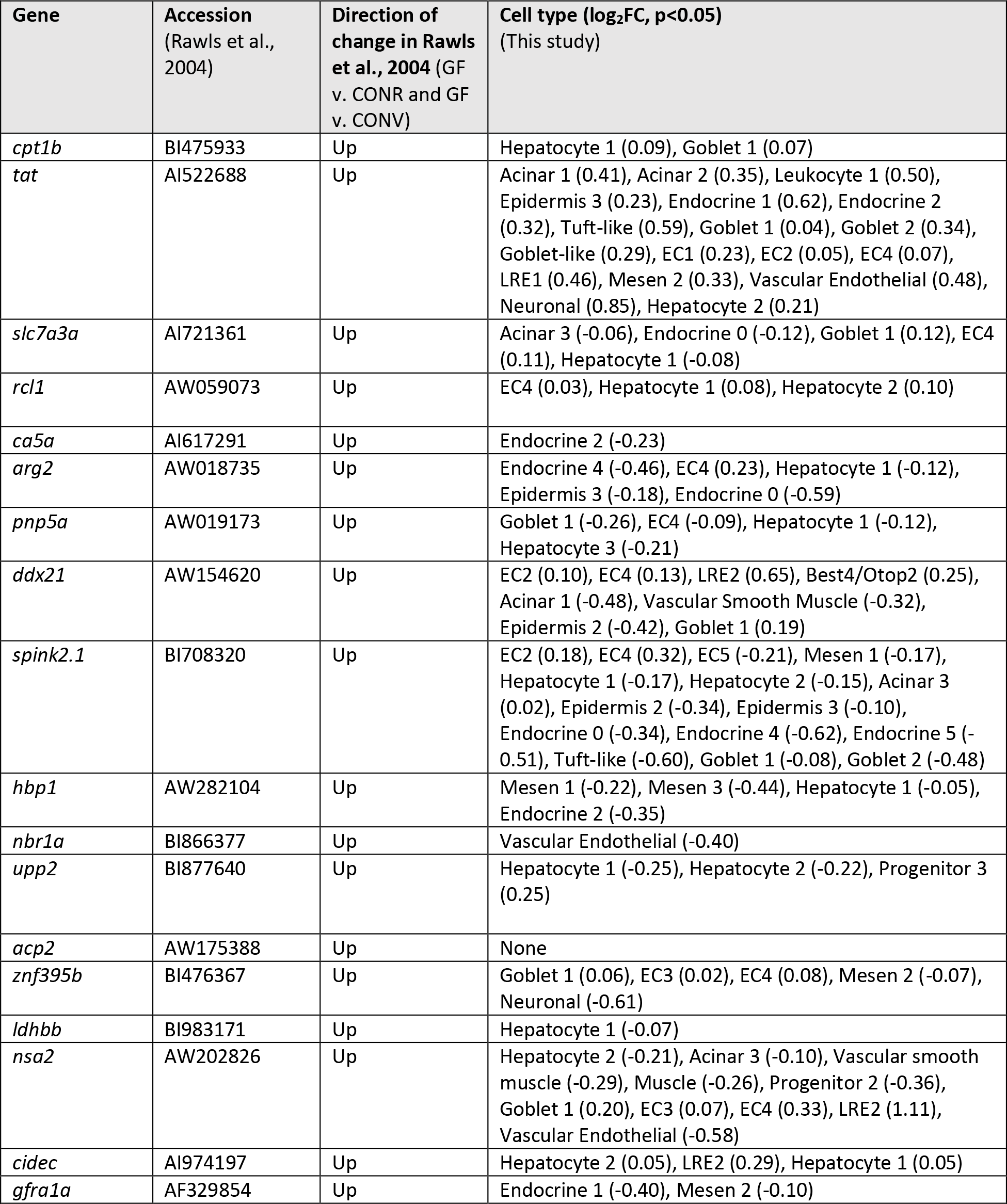
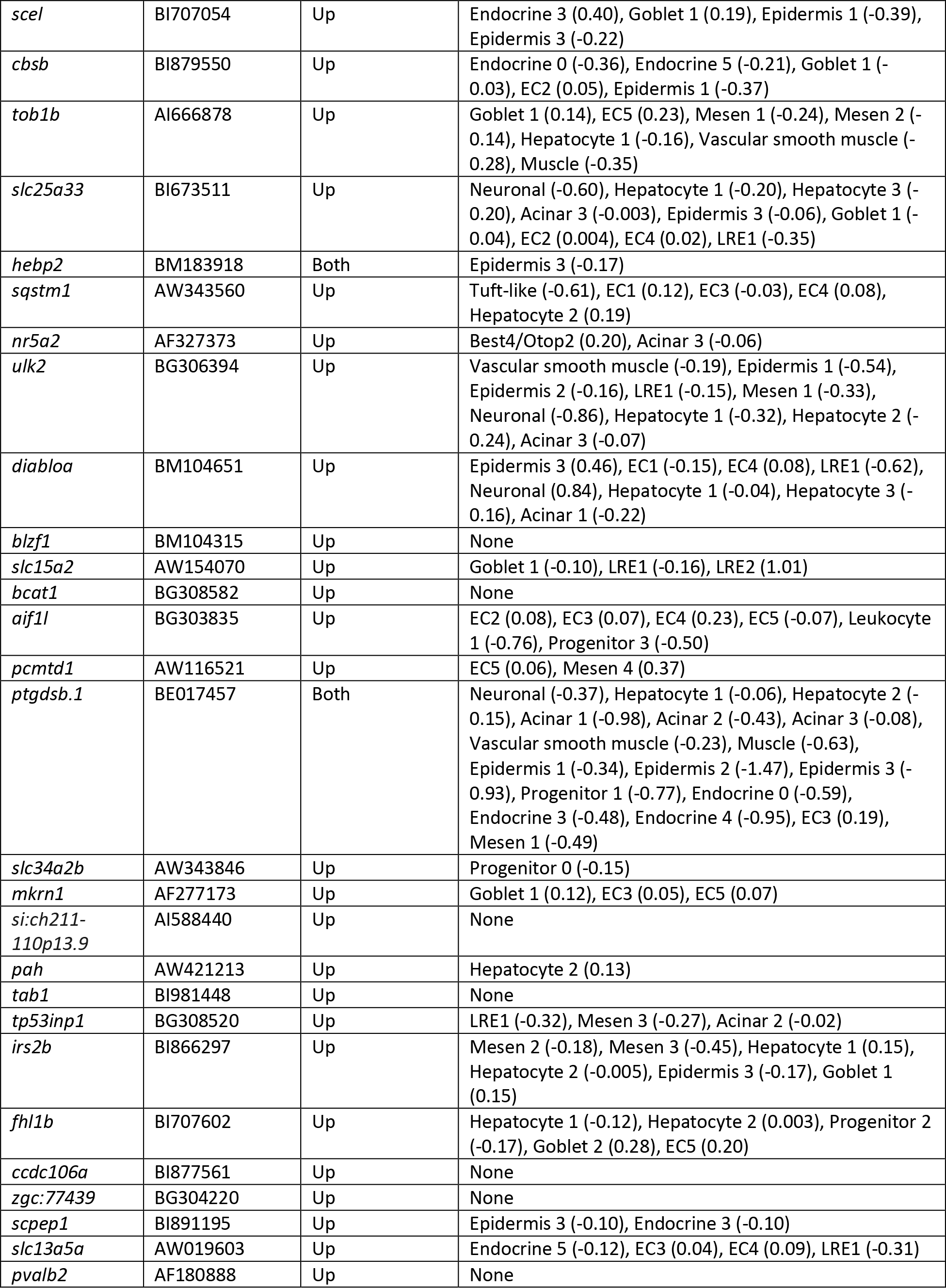
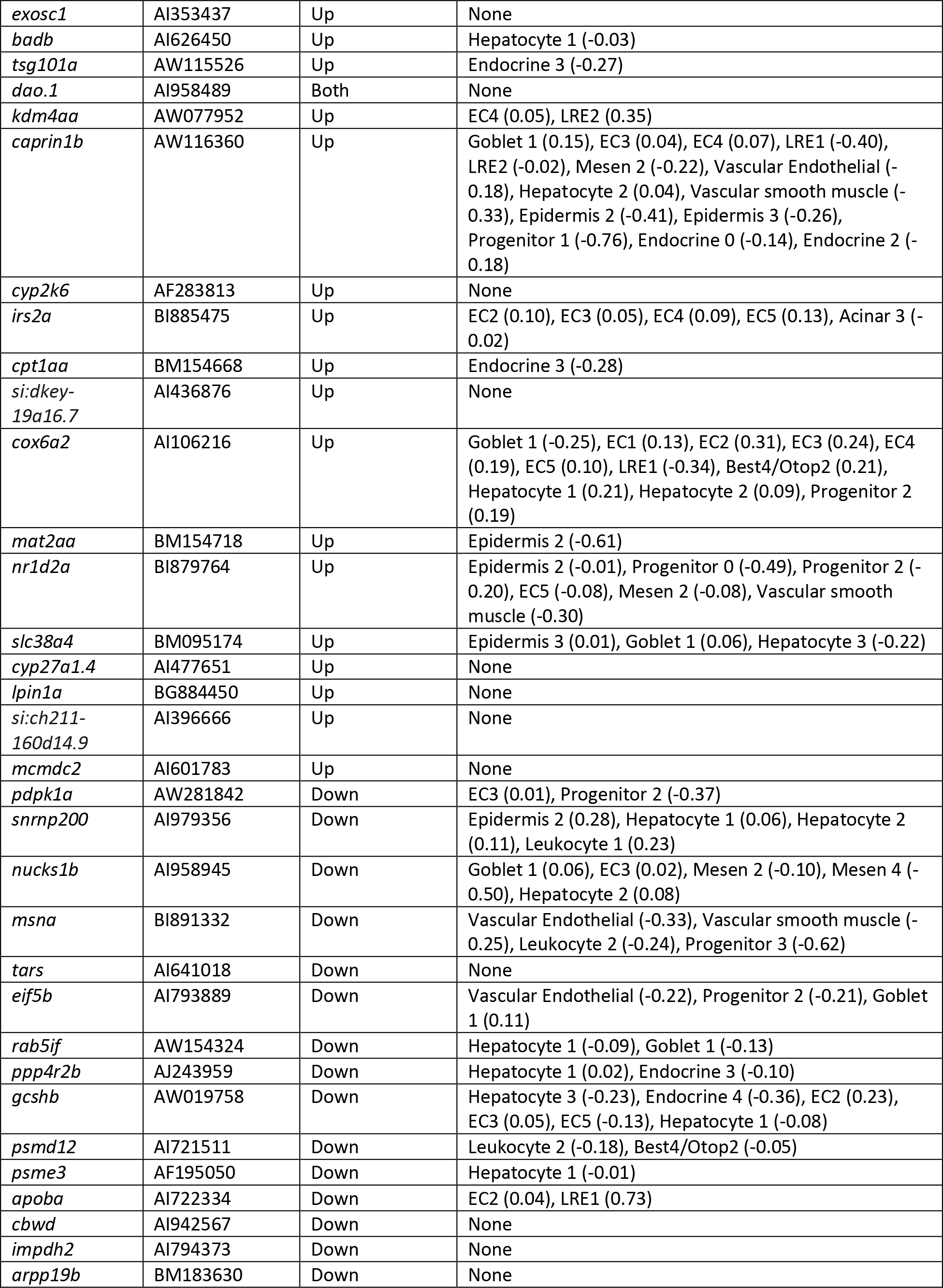
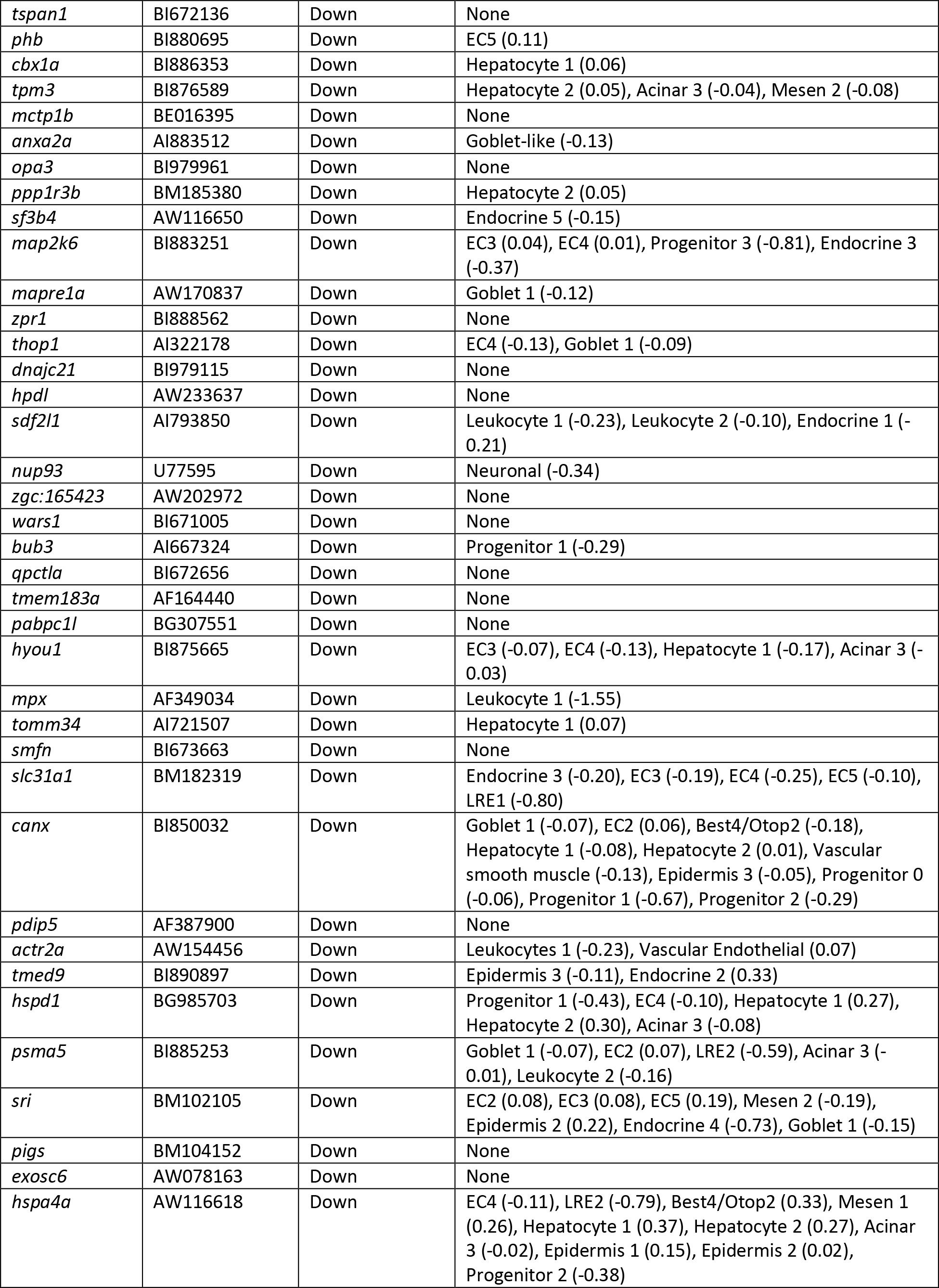
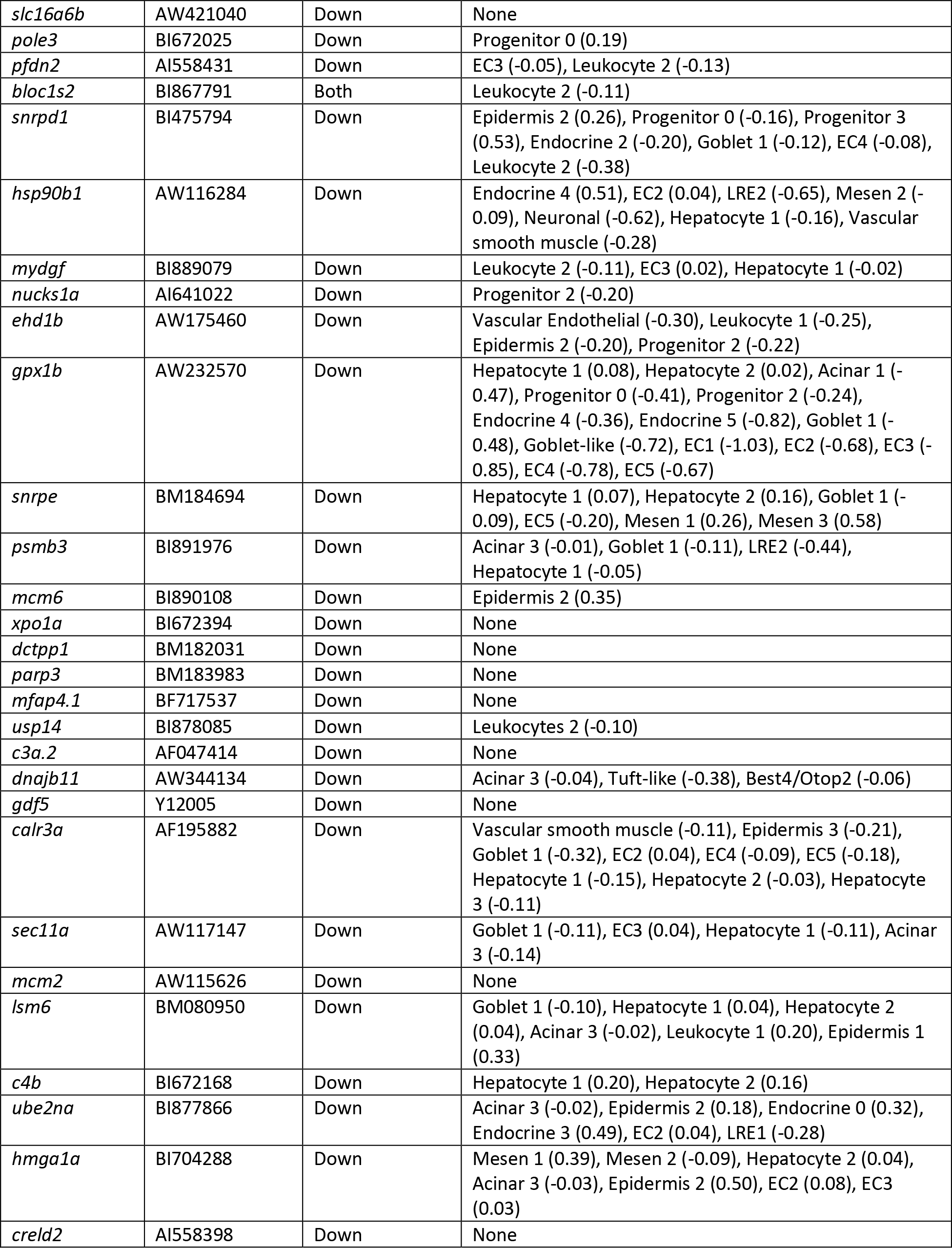
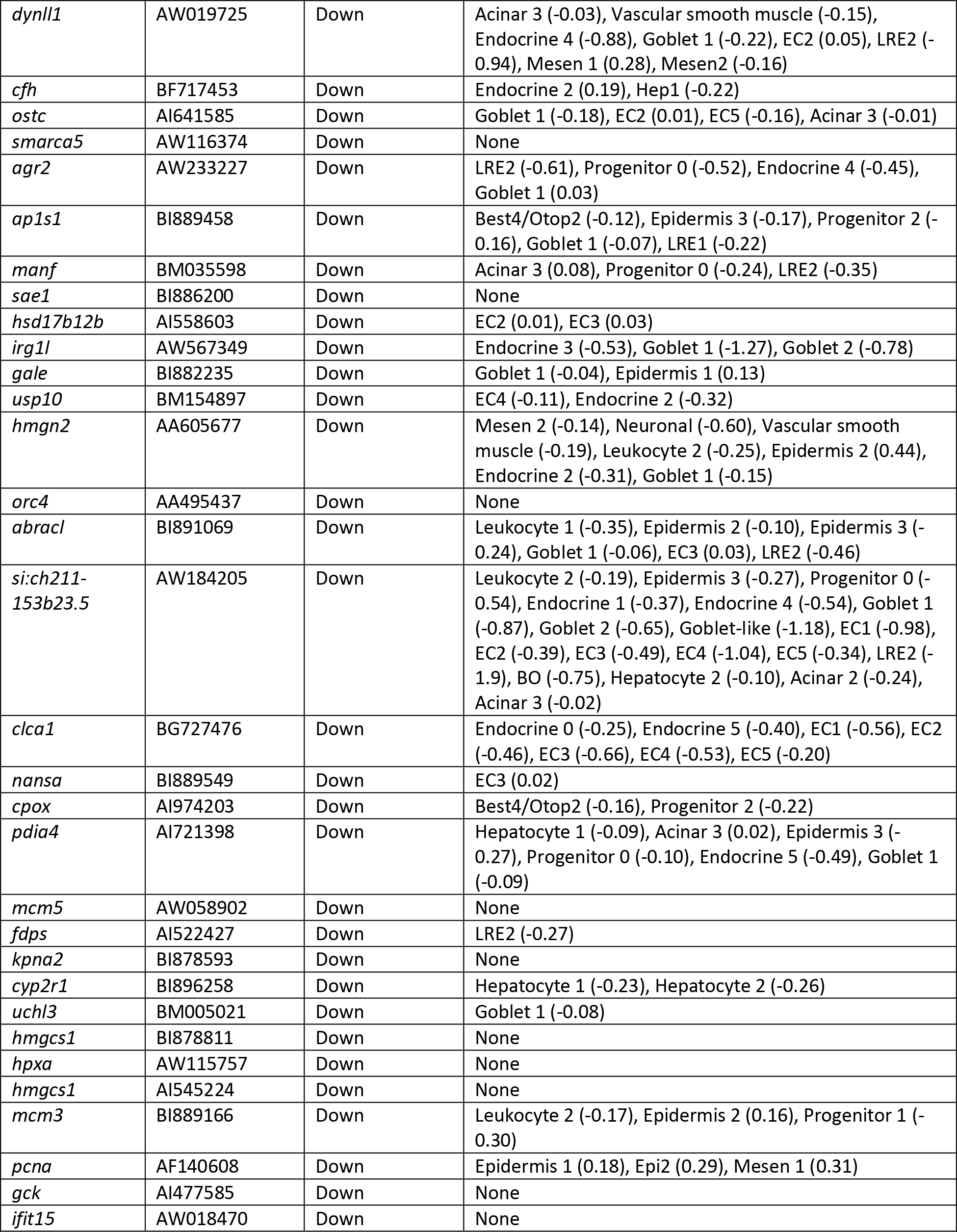
Microbe-responsive genes exhibit cell-specific changes upon bacterial colonization. Cell-type specific transcriptional changes are shown for genes whose expression changed in GF relative to both conventionally reared (CONR) and GF animals conventionalized at 3 dpf (CONV), as reported in Rawls et al., 2004.

**Figure S1.**
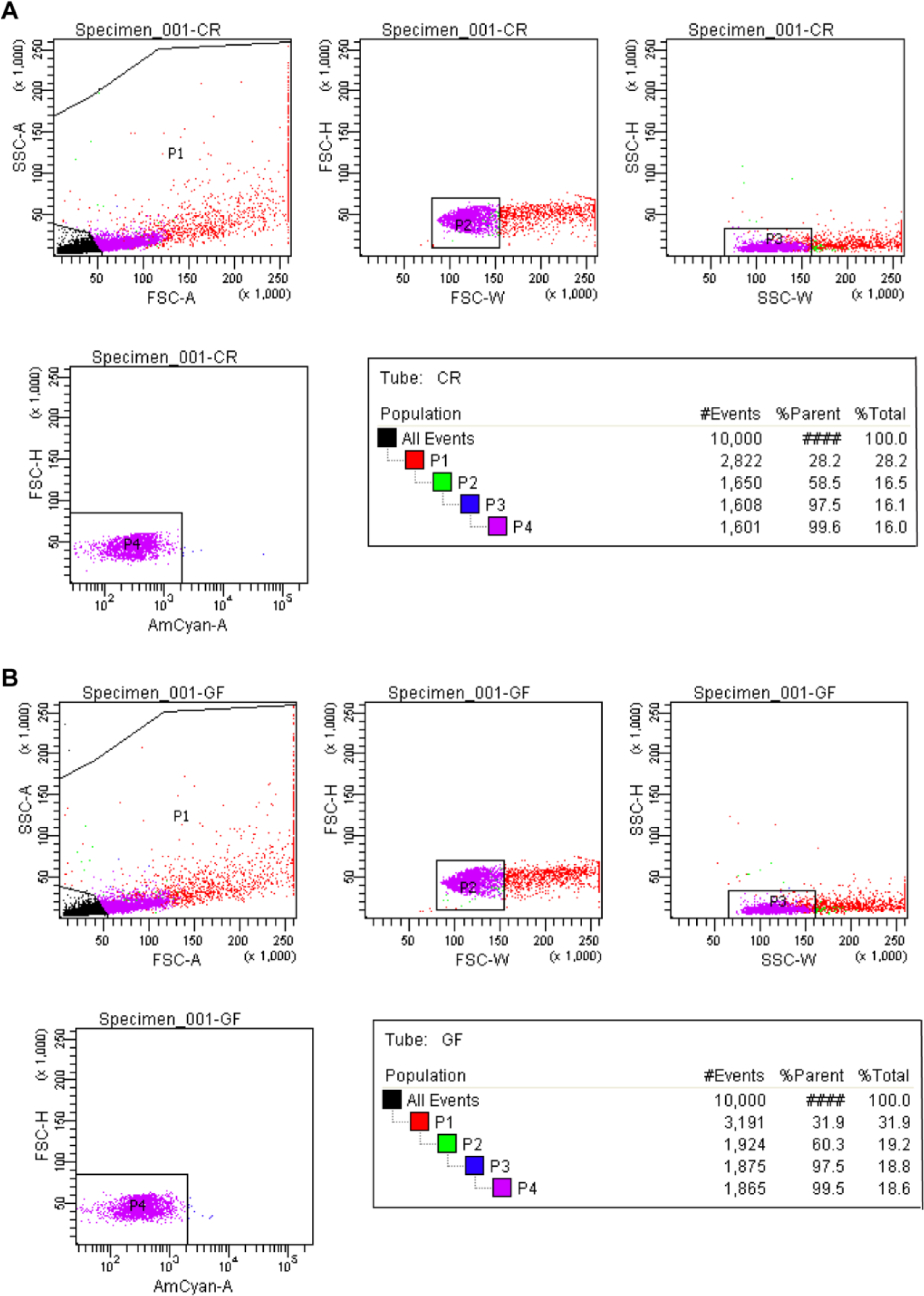
Isolation of single cells via fluorescence activated cell sorting. (*A-B*) Gating strategy for isolating dissociated single intestinal cells from CV (A) and GF (B) fish for replicate 1 of the single cell isolation protocol. Forward and side scatter were used to determine the single cell population, and Zombie Aqua viability die was used to select live cells.

**Figure S2.**
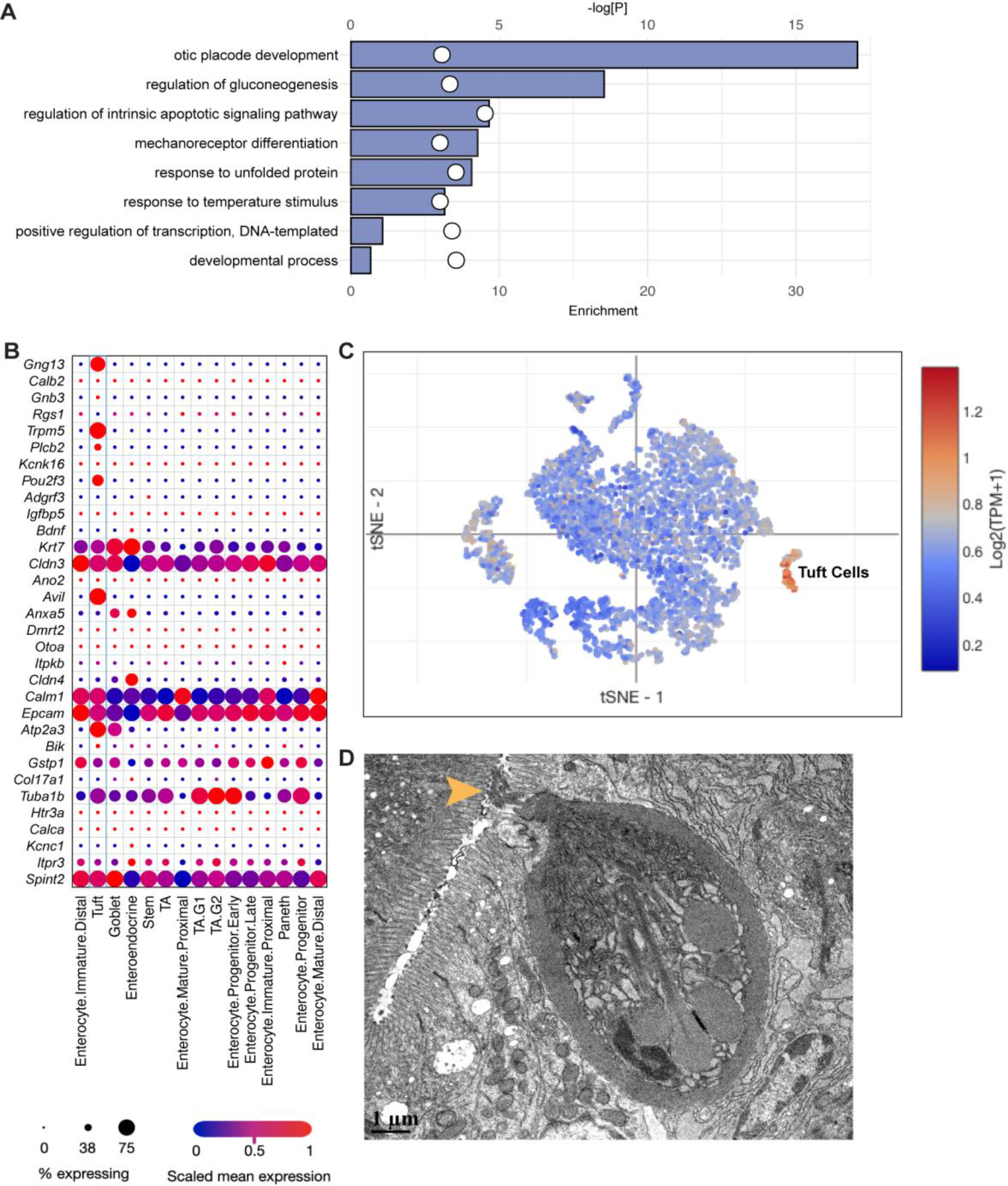
Identification of tuft-like cells in the zebrafish intestinal epithelium. (*A*) GO enrichment analysis of tuft-like cells, based on tuft-like cell genetic markers from the conventional single-cell RNA sequencing dataset. Top 8 GO terms are shown. Enrichment score is represented by bar length and p-value is indicated with white circles. (*B*-*C*) Expression of mouse orthologues for the top 50 zebrafish tuft-like cell marker genes in a single-cell RNA sequencing dataset of the mouse small intestinal epithelium from Haber et al., 2017, generated using the Broad Single Cell Portal. 32/50 genes had orthologues that were detected in the mouse small intestinal dataset. (*B*) Heatmap of mouse orthologue expression per mouse intestinal epithelial cell type, colored by scaled mean expression, where the size of the dot indicates proportion of expressing cells per cell type. (*C*) TSNE plot of mouse small intestinal epithelial cells, showing mean expression (Log2(TPM+1)) of 32 tuft-like marker gene orthologues per cell. Enrichment is evident in the annotated tuft cell clusters. (*D*) Transmission electron micrograph of the adult zebrafish posterior intestinal epithelium. Yellow arrowhead points to an apical tuft protruding through the epithelial brush border.

**Figure S3.**
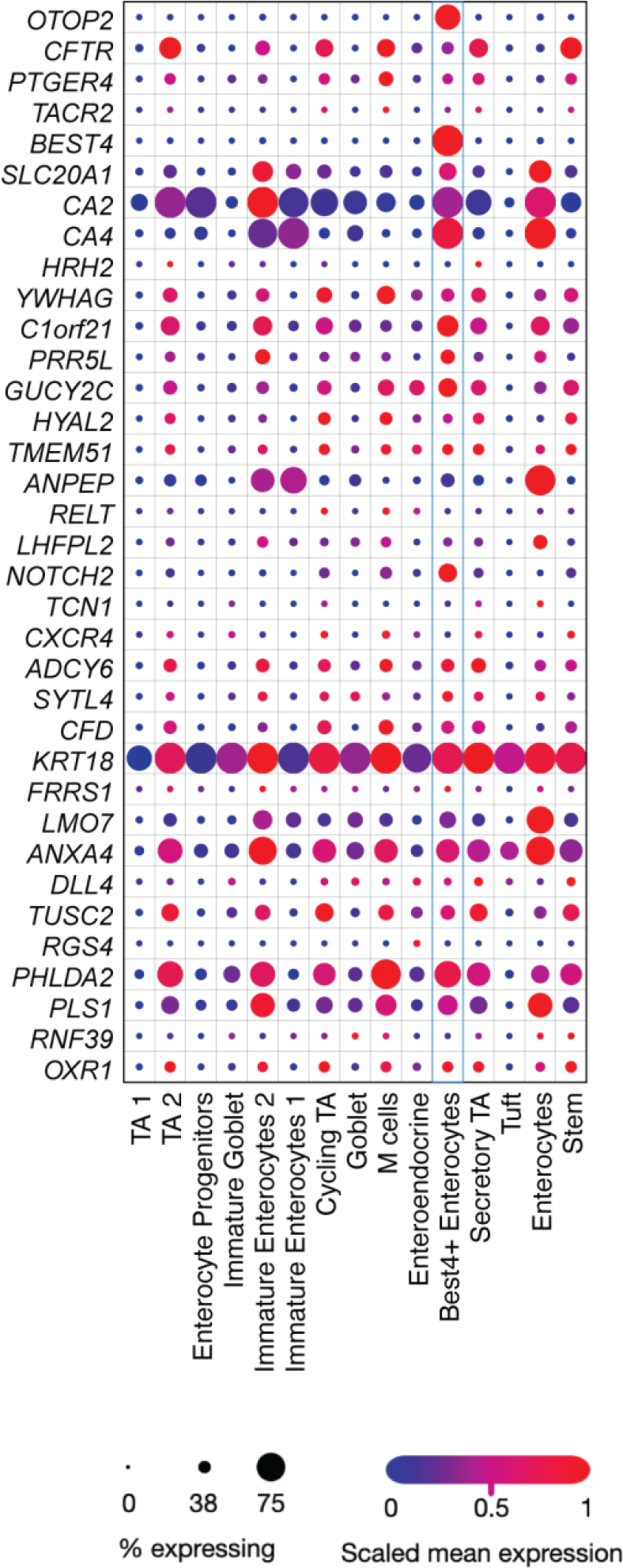
Analysis of zebrafish Best/Otop2 cell expression markers in the human colonic epithelium. Expression of human orthologues for the top 50 zebrafish Best4/Otop2 cell marker genes in a single-cell RNA sequencing dataset of the human colonic epithelium from Smillie et al., 2019, generated using the Broad Single Cell Portal (accession SCP259). 35/50 genes had orthologues that were detected in the human colonic dataset. Heatmap of human orthologue expression per epithelial cell type is shown, colored by scaled mean expression, where the size of the dot indicates proportion of expressing cells per cell type.

**Figure S4.**
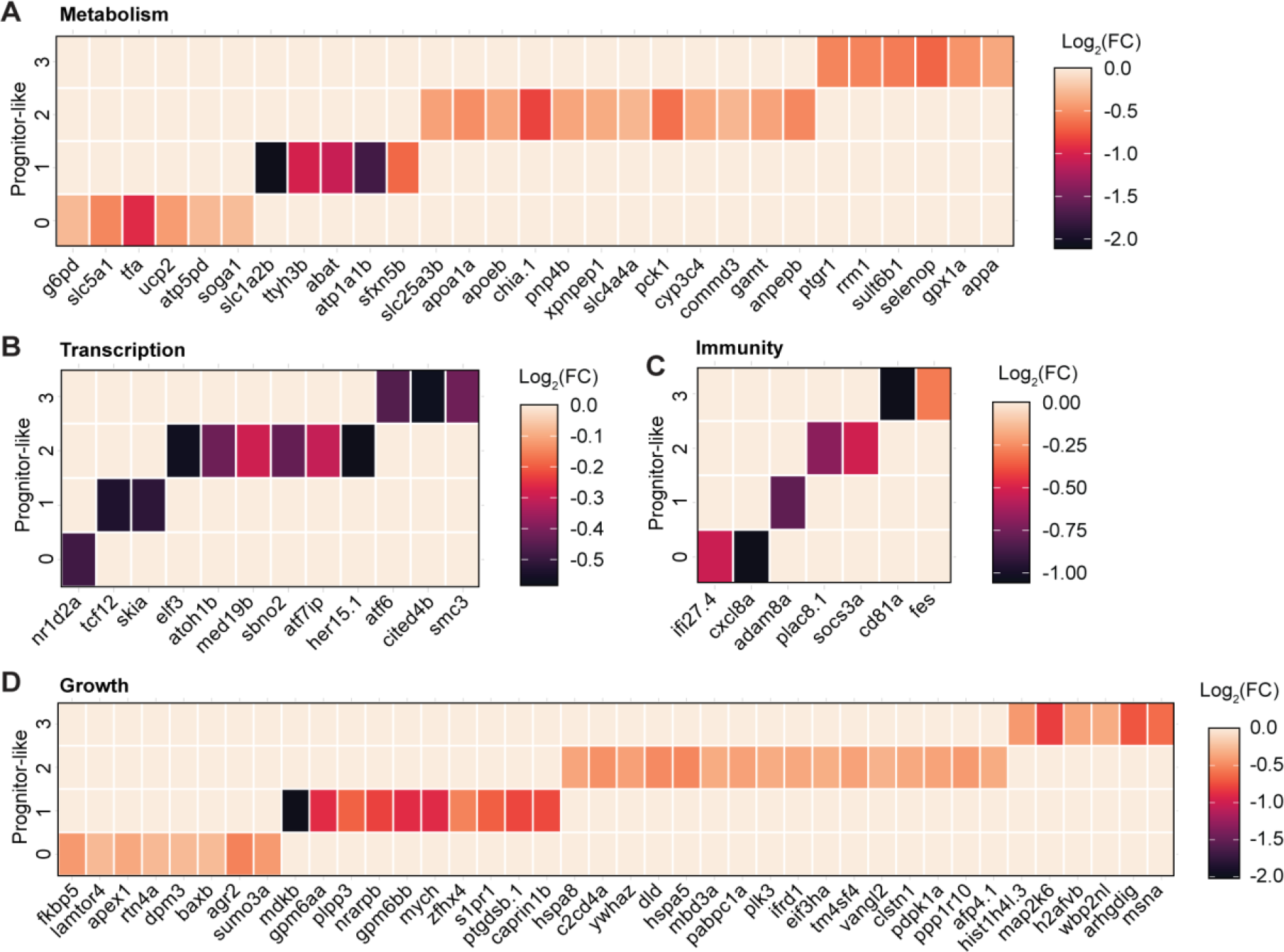
Microbes stimulate specialized processes in progenitor-like cell subsets. Heatmaps of differentially expressed genes (GF vs. CV, p<0.05) involved in metabolism (*A*), transcription (*B*), immunity (*C*) and growth (*D*), in progenitor-like subsets 0-3 (from Figure 1D-E), color coded according to Log2(FC). All non-zero value expression changes are significant (p<0.05) as determined with a non-parametric Wilcoxon rank sum test.

**Figure S5.**
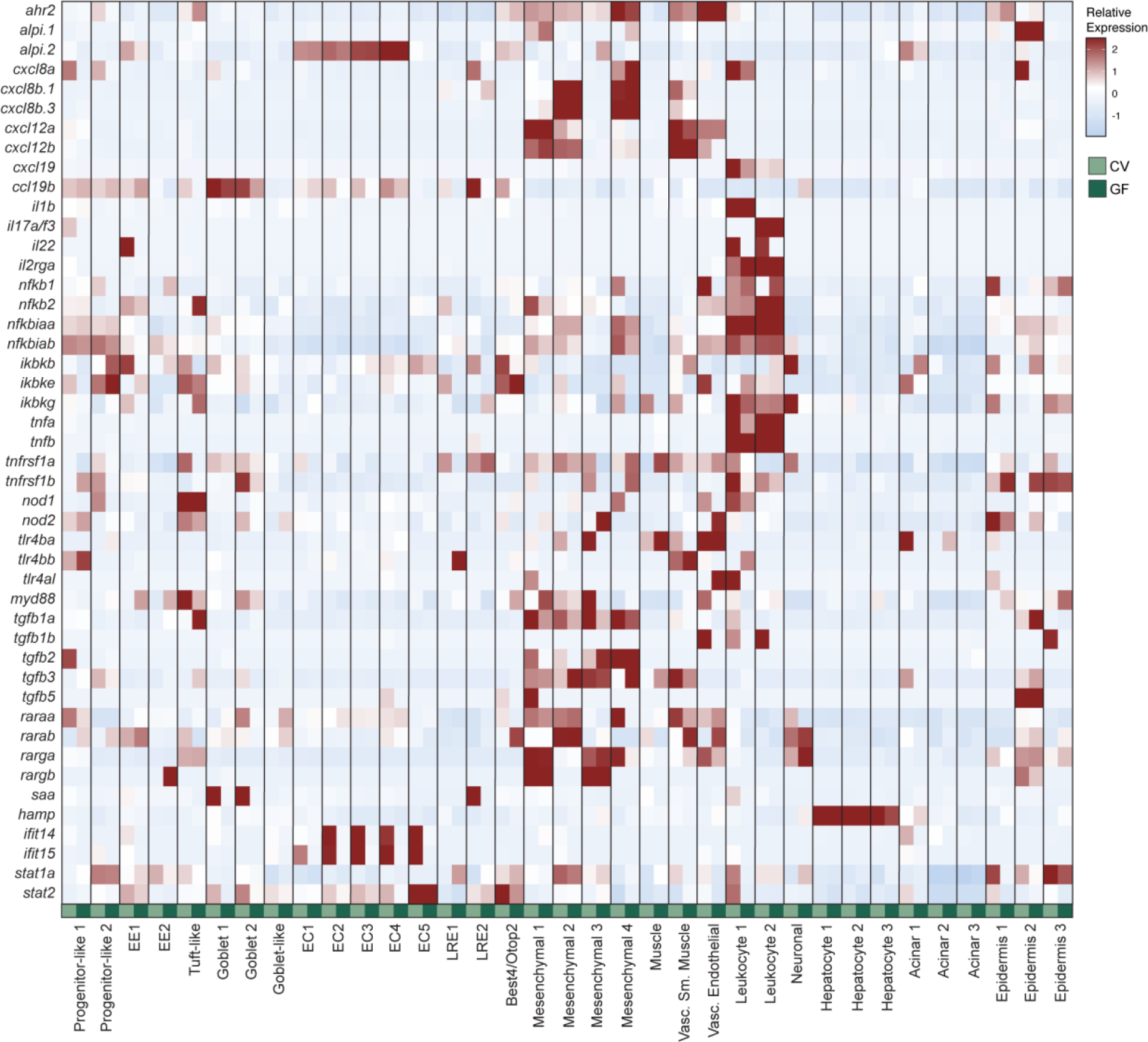
Immune gene expression across conventional and germ-free cell populations. Heatmap showing relative expression of a representative set of microbial sensors, NF-kB pathway components, cytokines and chemokines in CV and GF cell types. CV expression

**Figure S6.**
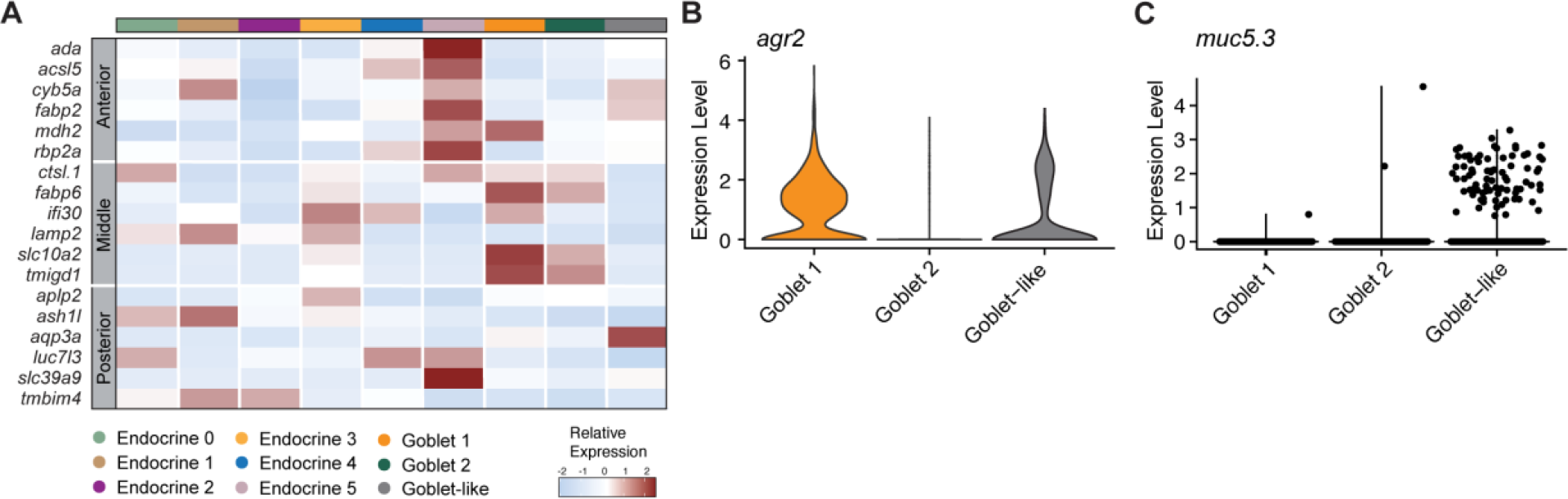
Secretory cell regional specification and goblet cell characterization. (*A*) Heatmap showing relative expression of established regional marker genes in each secretory cell type. (*B-C*) Violin plots for *agr2* (B) and *muc5.3* (C) expression in goblet and goblet-like clusters.

